# Detecting and Reversing Myocardial Ischemia Using an Artificially Intelligent Bioelectronic Medicine

**DOI:** 10.1101/2020.12.30.424900

**Authors:** PD Ganzer, MS Loeian, SR Roof, B Teng, L Lin, DA Friedenberg, IW Baumgart, EC Meyers, KS Chun, A Rich, WW Muir, DJ Weber, RL Hamlin

## Abstract

**Summary:** Myocardial ischemia is spontaneous, usually asymptomatic, and contributes to fatal cardiovascular consequences. Importantly, biological neural networks cannot reliably detect and correct myocardial ischemia on their own. In this study, we demonstrate an artificially intelligent and responsive bioelectronic medicine, where an artificial neural network (ANN) supplements biological neural networks enabling reliable detection and correction of myocardial ischemia. ANNs were first trained to decode spontaneous cardiovascular stress and myocardial ischemia with an overall accuracy of ∼92%. ANN-controlled vagus nerve stimulation (VNS) reversed the major biomarkers of myocardial ischemia with no side effects. In contrast, open-loop VNS or ANN-controlled VNS following a caudal vagotomy essentially failed to reverse correlates of myocardial ischemia. Lastly, variants of ANNs were used to meet clinically relevant needs, including interpretable visualizations and unsupervised detection of emerging cardiovascular stress states. Overall, these results demonstrate that ANNs can supplement deficient biological neural networks via an artificially intelligent bioelectronic medicine system.

## Introduction

Cardiovascular disease is responsible for a staggering ∼25-30% of mortality worldwide (World Health Organization, 2018). One prominent attribute of cardiovascular disease is myocardial ischemia – caused by a decrease in myocardial oxygen supply and / or an increase in myocardial oxygen demand (Ardehali &Ports, 1990; Deedwania &Carbajal, 1992; Hinderliter et al., 1991). Treating myocardial ischemia can reduce rates of myocardial injury, myocardial infarction, and death (Braun et al., 2018; Conti et al., 2012; Gutterman, 2009; Cohn, 1998). Unfortunately, treating myocardial ischemia is accompanied by several major challenges.

Roughly 75% of ischemic episodes are asymptomatic, where the heart can be irreversibly damaged without conscious awareness (Gutterman, 2009; Deedwania &Nelson, 1990; Rozanski &Berman, 1987; Cecchi et al., 1983). This significantly complicates the detection of myocardial ischemia, and clearly shows that biological neural networks are considerably deficient at detecting myocardial ischemia. Furthermore, myocardial ischemia can occur at random throughout the day (Schwartz et al., 2018; Gutterman, 2009; Cecchi et al., 1983), making it difficult to detect and treat. Supplementing deficient biological neural networks represents a promising approach to more effectively detect and potentially reverse myocardial ischemia.

In this study, we assessed the hypothesis that artificial neural networks (ANNs) can supplement deficient biological neural networks to detect, and even help correct, myocardial ischemia. To this end, we used an ANN that rapidly decodes events of spontaneous myocardial ischemia, and responsively triggers therapeutic closed-loop vagus nerve stimulation (VNS). Responsive closed-loop VNS may be an effective bioelectronic medicine for reversing ischemia mediated elevations in chronotropy, afterload, and myocardial oxygen demand (Capilupi et al., 2020; Levy &Schwartz, 1994; Ardell et al., 2015; Buck et al., 1981).

Although promising, implementing an ANN-controlled bioelectronic medicine for myocardial ischemia is difficult for several reasons. Events of myocardial ischemia are physiologically variable, within and across subjects (Patel et al., 1996; Celermajer et al., 1994; Deanfield &Spiegelhalter, 1990; Tzivoni et al., 1987). Furthermore, non-ischemic states have electrophysiological characteristics similar to myocardial ischemia (e.g., cardiac valve dysfunction, repolarization abnormalities, or an electrolyte imbalance; Michaelides et al., 2010; Sapin et al., 1991; Petrov et al., 2012; Gutterman, 2009). Therefore, detecting myocardial ischemia is complicated by significant biomarker variability and off-target states.

The majority of bioelectronic medicines use preprogrammed open-loop stimulation schedules. However, a closed-loop bioelectronic medicine that selectively responds when needed can optimize therapeutic efficacy (Wright et al., 2016; Sun &Morell, 2014; Hays, 2016; Ganzer et al., 2018; Ganzer &Sharma, 2019). Also, myocardial ischemia occurs throughout the day at random (Schwartz et al., 2018; Gutterman, 2009; Cecchi et al., 1983). Therefore, an effective bioelectronic treatment for myocardial ischemia may need to leverage responsive closed-loop control – where on-demand stimulation is autonomously triggered when needed for benefit.

Lastly, artificial intelligence (AI) is becoming a powerful tool in medicine. Importantly, AI enabled medicines must be easily interpretable for widespread adoption (Vellido, 2019; Tonekaboni et al., 2019; Tjoa, E., &Guan, 2019). AI interpretability can be enhanced using visualizations. However, it can be challenging to create interpretable visualizations of both high-dimensional data and complex algorithm decisions (Vellido, 2019, 2012, &2011; Liu et al., 2017; Zahavy et al., 2016). Furthermore, disease pathophysiology and biomarker data are always changing – over time, subjects can experience new forms of cardiovascular stress, and new pathophysiological states may emerge (Epel et al., 2018; Schwartz et al., 2018). Therefore, future AI enabled medicines will need to be both interpretable and adaptive to physiological changes.

## Results

### Inducing Acute Myocardial Ischemia

Clinical myocardial ischemia is associated with enhanced catecholamine tone, increased afterload, and changes to the myocardial oxygen supply / demand ratio commonly lasting 30 seconds or more (Gutterman, 2009; Rocco et al., 1986; Rehman et al., 1997; Hinderliter et al., 1991; Deedwania &Nelson, 1990). We modeled these attributes of clinical myocardial ischemia using injections of dobutamine and norepinephrine in rats. Dobutamine (primarily a β1 receptor agonist) is commonly used in the clinic to induce cardiovascular stress and subsequent myocardial oxygen demand (Mandapaka &Hundley, 2006). Norepinephrine (primarily an α1 receptor agonist) is extensively implicated in both coronary and peripheral vasoconstriction, cardiovascular stress, and myocardial ischemia (Heusch G &Ross, 1990; Kawada et al., 2002). Our approach was motivated by previous studies modeling cardiovascular stress and acute myocardial ischemia using injected catecholamines (Vimercati et al., 2012; Segar et al., 1995; Barger et al., 1961; Lepeschkin et al., 1960). We used three types of injections (schematic of experimental interfaces: Supplemental Fig. S1A): 1) dobutamine alone (D), 2) norepinephrine alone (NE), or dobutamine and norepinephrine combined (D+NE). Injection protocols consisted of an initial rest period followed by a 2-minute injection period (injection start = vertical dashed line, Fig. 1B-1E).

**Figure 1.**
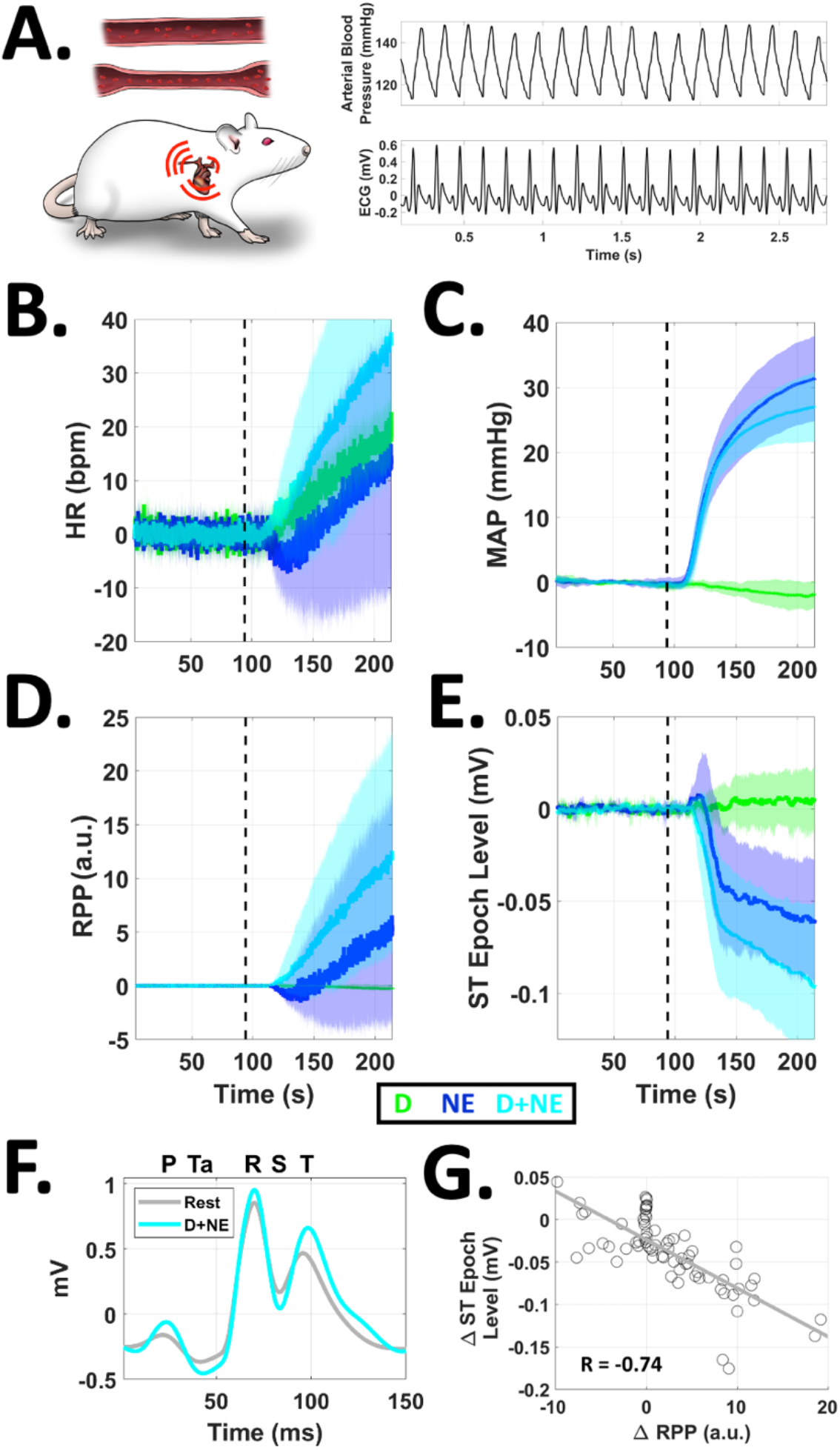
Inducing Acute Myocardial Ischemia *In Vivo*. **A.** Cartoon of rat experiments for inducing cardiovascular stress and myocardial ischemia (blood vessel / vasoconstriction cartoon: top left; heart / tachycardia cartoon: bottom left; arterial blood pressure waveforms: top right; electrocardiogram or ECG waveforms: bottom right). Heart rate (HR, **B**), mean arterial pressure (MAP, **C**), rate-pressure product (RPP, **D**), and ST epoch level (**E**) were differentially modulated following a dobutamine (D, green), norepinephrine (NE, blue), or combined dobutamine and norepinephrine (D+NE, cyan) injection, indicative of cardiovascular stress and myocardial ischemia (time series include lighter shaded regions = ± 95% confidence intervals; vertical black dashed line = time of injection). **F.** Representative ECG during rest (gray) or D+NE induced myocardial ischemia (cyan). Note the pronounced suppression of ECG epochs during both diastole and systole, indicative of ischemic currents (ECG waveforms respectively averaged across 10 seconds from each given period; relative P, Ta, R, S, and T ECG wave time points shown at top of the panel). **G.** ST epoch level depression was significantly correlated with RPP (RPP = an index of myocardial oxygen consumption). These results demonstrate that injected catecholamines differentially impact cardiovascular states and can induce acute myocardial ischemia.

Each injection type differentially impacted traditional biomarkers of cardiovascular stress and myocardial ischemia, including heart rate (Fig. 1B), mean arterial pressure (MAP, Fig. 1C), rate-pressure product (RPP, Fig. 1D), and ST epoch level (Fig. 1E). The combined D+NE injection tended to have a larger effect on two biomarkers intimately related to myocardial ischemia: 1) RPP, an index of myocardial oxygen consumption (Gobel et al., 1978; Detry et al., 1970); and 2) ST epoch depression, a classic electrophysiological correlate of subendocardial ischemia (Klabunde, 2017). A representative averaged ECG is shown before (Rest, gray; Fig. 1F) and during D+NE induced myocardial ischemia (D+NE, cyan; Fig. 1F). Electrophysiological correlates of decreased myocardial membrane potential and ischemic currents (Klabunde, 2017; Janse, 2007; Cinca et al., 1980; Kleber et al., 1978) were observed during both systole (during QRS) and diastole (during the ST and Ta epochs). Specifically, D+NE induced a maximal ST epoch depression up to ∼0.1 mV. ST epoch depression was significantly correlated to RPP across all injections (Fig. 1G: R = -0.74; p<0.001). Therefore, increased subendocardial ischemia was associated with higher myocardial oxygen consumption. These results demonstrate that catecholamine injections induce correlates of cardiovascular stress and acute myocardial ischemia.

### Creating Features for Cardiovascular State Decoding

We next created a broader set of features from the ECG and blood pressure signals for state decoding and target ischemia detection (ECG feature schematic: Fig. 2A; blood pressure feature schematic: Fig. 2B). The broader 13 element feature vector further quantified several biomarkers, such as ECG segment durations (ms), relative ECG wave point levels (mV), blood pressures during diastole and systole (mmHg), and breath rate. Changes in features were assessed for D, NE, and D+NE, quantified with respect to baseline levels (i.e., Δ relative to baseline).

**Figure 2.**
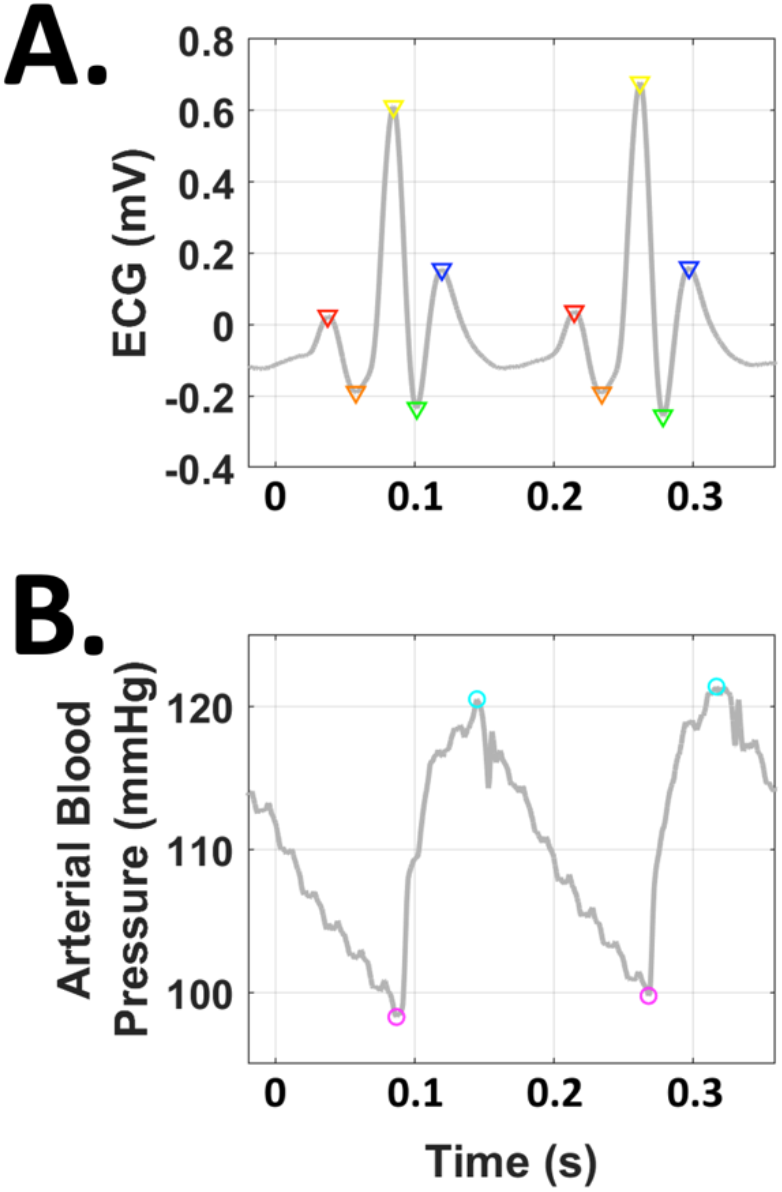
Schematic of Cardiovascular Feature Components. All 13 cardiovascular features (shown in Figure 3) were derived from the ECG (**A**) or arterial blood pressure (**B**) signals. Across wave cycles, we identified correlates of the P wave (red triangles, atrial depolarization), Ta wave (orange triangles, atrial repolarization), R wave (yellow triangles, ventricular depolarization), S wave (green triangles, nadir between ventricular depolarization and repolarization), T wave (blue triangles, ventricular repolarization), diastolic pressure (magenta circles), and systolic pressure (cyan circles). Breath rate was derived from the linear envelope of the blood pressure signal. The 13-element feature vector was calculated every 100 ms, and averaged over a 4 s sliding window. Please see the *On-Line Cardiovascular Signal Conditioning and Feature Extraction* section of the methods for more details on feature extraction.

Additional biomarkers of cardiovascular stress and myocardial ischemia were observed across the added features (Klabunde, 2017; Janse, 2007; Rehman et al., 1997; Deedwania &Nelson, 1990; Cinca et al., 1980; Kleber et al., 1978), including increases in pulse pressure, decreases in myocardial conduction velocity, and depression of other ECG wave points indicative of ischemic currents (Fig. 3A: ECG features; Fig. 3B: blood pressure features; Fig. 3C, pulmonary feature; Table 1: omnibus ANOVA results). The combined injection of D+NE also had a maximal effect on this broader set of features compared to D or NE. These results demonstrate the more distributed impact of catecholamines on broader features of cardiovascular stress and myocardial ischemia.

**Figure 3.**
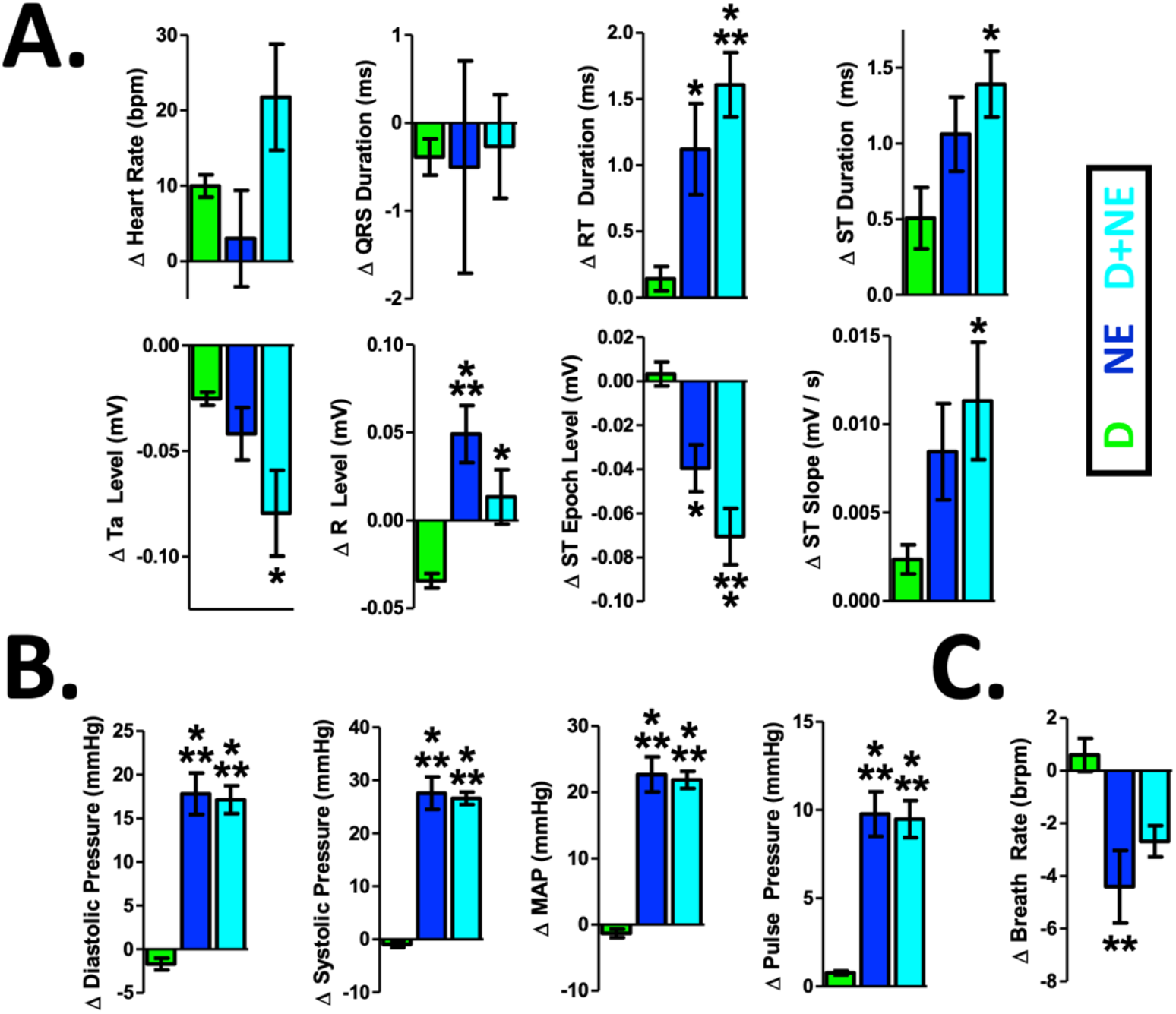
Cardiovascular Feature Changes During Induction of Cardiovascular Stress and Myocardial Ischemia. A more detailed 13 element cardiovascular feature vector was created for eventual decoding of cardiovascular state. Dobutamine (D, green), norepinephrine (NE, blue), or a combination of dobutamine and norepinephrine (D+NE, cyan) induced significant changes to features related to the ECG (**A**), blood pressure (**B**), and pulmonary function (**C**). Of note, D+NE maximally impacted several cardiovascular features (* = different from D at p<0.05; ** = different from D at p<0.01; *** = different from D at p<0.001). This 13-element feature vector was next used for decoding cardiovascular stress and myocardial ischemia. Data presented are mean ± SEM.

**Table 1:** Omnibus ANOVA results (related to Fig. 3).

|  | **Omnibus ANOVA Results** |
| --- | --- |
| **$\Delta$ Feature 1 (Heart Rate)** | $F[2,24] = 2.8, p = 0.07$ |
| **$\Delta$ Feature 2 (QRS Duration)** | $F[2,24] = 0.02, p = 0.97$ |
| **$\Delta$ Feature 3 (RT Duration)** | $F[2,24] = 8.9, p < 0.01$ |
| **$\Delta$ Feature 4 (ST Duration)** | $F[2,24] = 4, p < 0.05$ |
| **$\Delta$ Feature 5 (Ta Level)** | $F[2,24] = 4, p < 0.05$ |
| **$\Delta$ Feature 6 (R Level)** | $F[2,24] = 10.1, p < 0.001$ |
| **$\Delta$ Feature 7 (ST Epoch Level)** | $F[2,24] = 13.3, p < 0.001$ |
| **$\Delta$ Feature 8 (ST Slope)** | $F[2,24] = 3.2, p < 0.05$ |
| **$\Delta$ Feature 9 (Diastolic Pressure)** | $F[2,24] = 44.2, p < 0.001$ |
| **$\Delta$ Feature 10 (Systolic Pressure)** | $F[2,24] = 71.3, p < 0.001$ |
| **$\Delta$ Feature 11 (Mean Arterial Pressure)** | $F[2,24] = 60.7, p < 0.001$ |
| **$\Delta$ Feature 12 (Pulse Pressure)** | $F[2,24] = 28.9, p < 0.001$ |
| **$\Delta$ Feature 13 (Breath Rate)** | $F[2,24] = 7.3, p < 0.01$ |

Importantly, biomarker features of myocardial ischemia can be variable within and across subjects (Patel et al., 1996; Celermajer et al., 1994; Deanfield &Spiegelhalter, 1990; Tzivoni et al., 1987). Furthermore, seemingly separate cardiovascular states can exhibit highly correlated biomarkers and thus statistically overlap (Sharma &Gedeon, 2012; Michaelides et al., 2010; Petrov et al., 2012; Sapin et al., 1991; Gutterman, 2009). Therefore, both biomarker variability and correlation across states should be attributes of a myocardial ischemia model.

The cardiovascular feature data (from Fig. 3) exhibited variability and disorder comparable to human cardiovascular data recorded in either the intensive care unit (Supplemental Fig. S2A; Kim et al., 2016; Goldberger et al., 2000) or during ambulatory episodes of myocardial ischemia (Supplemental Fig. S2B; Taddei et al., 1992; Jager et al., 2003; Goldberger et al., 2000). Furthermore, there was a significant correlation between NE and D+NE, even though they are distinct and separate cardiovascular stress states (Supplemental Fig. S2C). These findings demonstrate that the recorded cardiovascular features importantly model the variability and state overlap seen during human cardiovascular stress and myocardial ischemia, a clinically relevant challenge for cardiovascular state decoding.

### Decoding Complex Cardiovascular States Using an Artificial Neural Network

Biological neural networks are largely incapable of detecting myocardial ischemia (∼75% of episodes are asymptomatic: Gutterman, 2009; Deedwania &Nelson, 1990; Rozanski &Berman, 1987; Cecchi et al., 1983). An artificial neural network (ANN) may be able to supplement deficient biological neural networks to reliably detect, and even help correct, myocardial ischemia. We developed an ANN architecture to decode cardiovascular states during cardiovascular stress and myocardial ischemia, comprised of both a hidden dense layer and a hidden long short-term memory (LSTM) layer (4 total layers, schematic in Supplemental Fig. S3A; see the *Decoding Myocardial Demand Ischemia Using an Artificial Neural Network* section of the methods for more details). A LSTM layer was incorporated to detect long-term dependencies across time in the cardiovascular data and potentially enhance decoding performance (Murat et al., 2020; Gers et al., 1999). The output of the ANN is a continuous prediction score across the 4 states: rest (no drug injected), D, NE, or D+NE (example decoder outputs during a D+NE injection: Supplemental Fig. S3B).

Despite significant feature variability and state overlap, the ANN decoded cardiovascular state with high overall accuracy (∼92%, Fig. 4 &Supplemental Fig. S3C; F[3,36] = 163.5, p < 0.001). Replacing the LSTM layer with a normal dense layer removed the network’s ability to assess long term dependencies in the signal, significantly decreasing accuracy (i.e., an ANN-NO-LSTM architecture; Supplemental Fig. S3C). The ANN also outperformed other classifiers such as a support vector machine or a linear discriminant analysis (Supplemental Fig. S3C).

**Figure 4.**
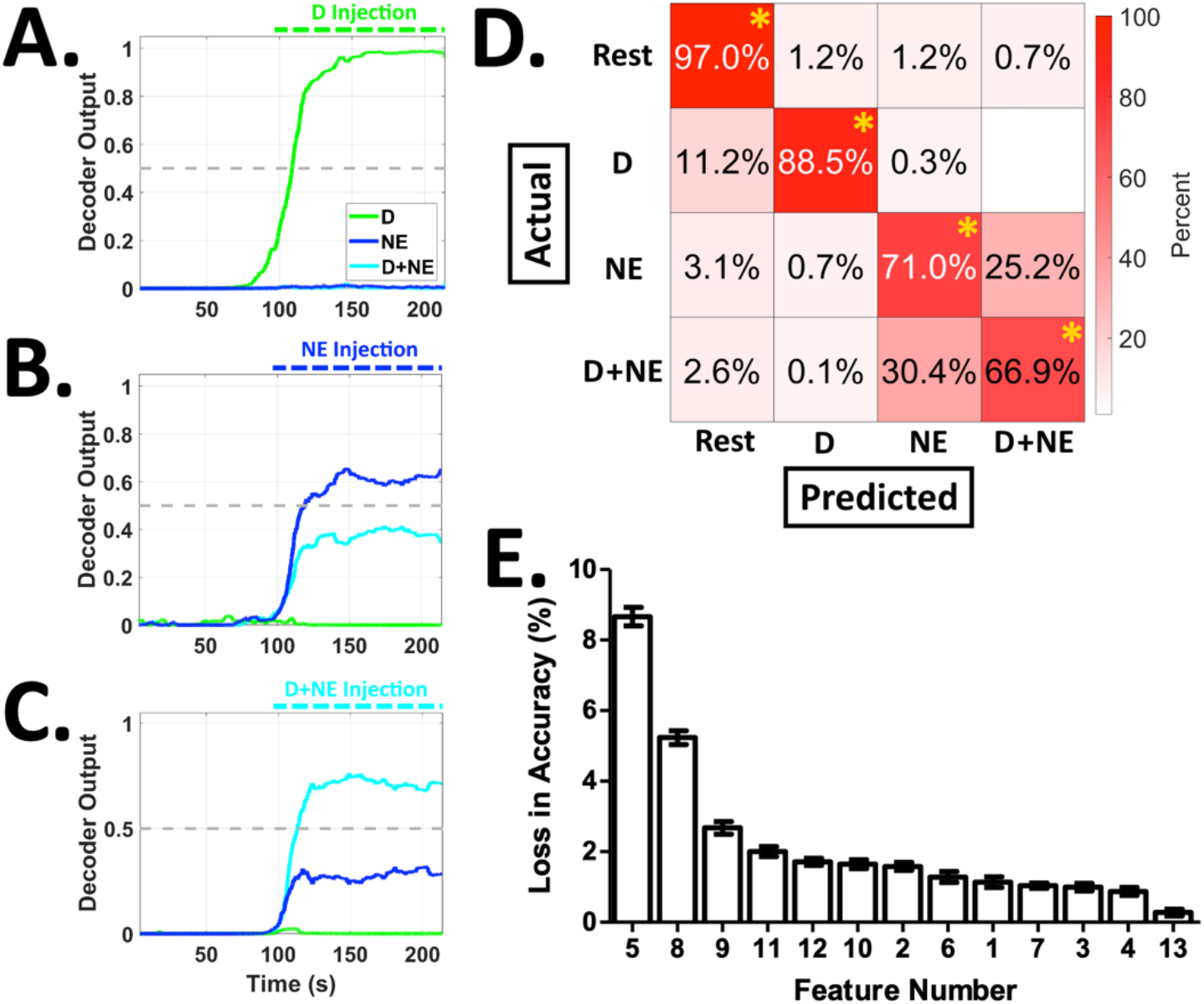
The ANN Accurately Classifies Cardiovascular Stress States and is Significantly Impacted by the Removal of Features Related to Cardiac Electrophysiology &Vascular Resistance. The ANN was challenged to decode complex cardiovascular state changes (cardiovascular feature variability and state overlap assessment: Supplemental Fig. S2) across a total of 4 classes: Rest, D, NE, or D+NE. Continuous decoder outputs are shown for the injection of D (panel **A**), NE (panel **B**), or D+NE (panel **C**). The ANN performed with a high overall accuracy (∼92%) and sensitivity (∼86%) (confusion matrix showing average performance values: panel **D;** * = above chance at p<0.001). **E.** The removal of features related to ECG ischemic currents (features 5 &8) or blood pressure (features 9-12) led to the largest losses in decoding accuracy. These results show that an ANN can accurately decode cardiovascular states, despite significant cardiovascular feature variability and state overlap. Data presented are mean ± SEM.

The ANN exhibited an overall sensitivity of ∼86% (example decoder outputs: Fig. 4A-4C; confusion matrix showing average performance values: Fig. 4D). Although the ANN had an overall accuracy of ∼92%, the most common misclassification was between the NE and D+NE classes (potentially due to their high degree of variability and correlation, as shown in Fig. 1, Fig. 3, and Supplemental Fig. S2C). Features related to ECG wave point depression, ischemic currents, and vascular resistance were the most important features for ANN decoding performance (Fig. 4E and Table 2). Lastly, a fixed ANN trained on subsets of the data robustly generalized to testing days spread out over several months and different animals (Supplemental Fig. S4). Overall, these results show that ANNs can robustly decode complex cardiovascular states to supplement deficient biological neural networks.

**Table 2.** Feature number and name, ranked according to loss in accuracy (i.e., highest loss to lowest loss; related to Fig. 4E)

| Feature Number (Fig. 4E) | 5 | 8 | 9 | 11 | 12 | 10 | 2 | 6 | 1 | 7 | 3 | 4 | 13 |
| --- | --- | --- | --- | --- | --- | --- | --- | --- | --- | --- | --- | --- | --- |
| Feature Name | Ta Level | ST Slope | Diastolic Pressure | Mean Arterial Pressure | Pulse Pressure | Systolic Pressure | QRS Duration | R Level | Heart Rate | ST Epoch Level | RT Duration | ST Length | Breath Rate |

### Responsive ANN Controlled Vagus Nerve Stimulation Reverses Myocardial Ischemia

Myocardial ischemia can cause irreversible heart damage if not treated rapidly. Therefore, beyond rapid detection alone, rapid myocardial ischemia correction is also needed. We next leveraged the ANN decoder to enable closed-loop vagus nerve stimulation (VNS) and potentially reverse myocardial ischemia (i.e., ANN-VNS; cartoon schematic: Fig. 5A). VNS can decrease chronotropy, afterload, and myocardial oxygen demand (Capilupi et al., 2020; Levy &Schwartz, 1994; Buck et al., 1981), all factors that are elevated during spontaneous myocardial ischemia (Svensson et al., 2001; Rehman et al., 1997; Deedwania &Carbajal, 1992; Hinderliter et al., 1991; Deedwania &Nelson, 1990). D+NE targets catecholamine receptors relevant for myocardial ischemia and has a maximal effect on the recorded features. Therefore, we targeted D+NE induced myocardial ischemia for detection and correction, using ANN-VNS.

**Figure 5.**
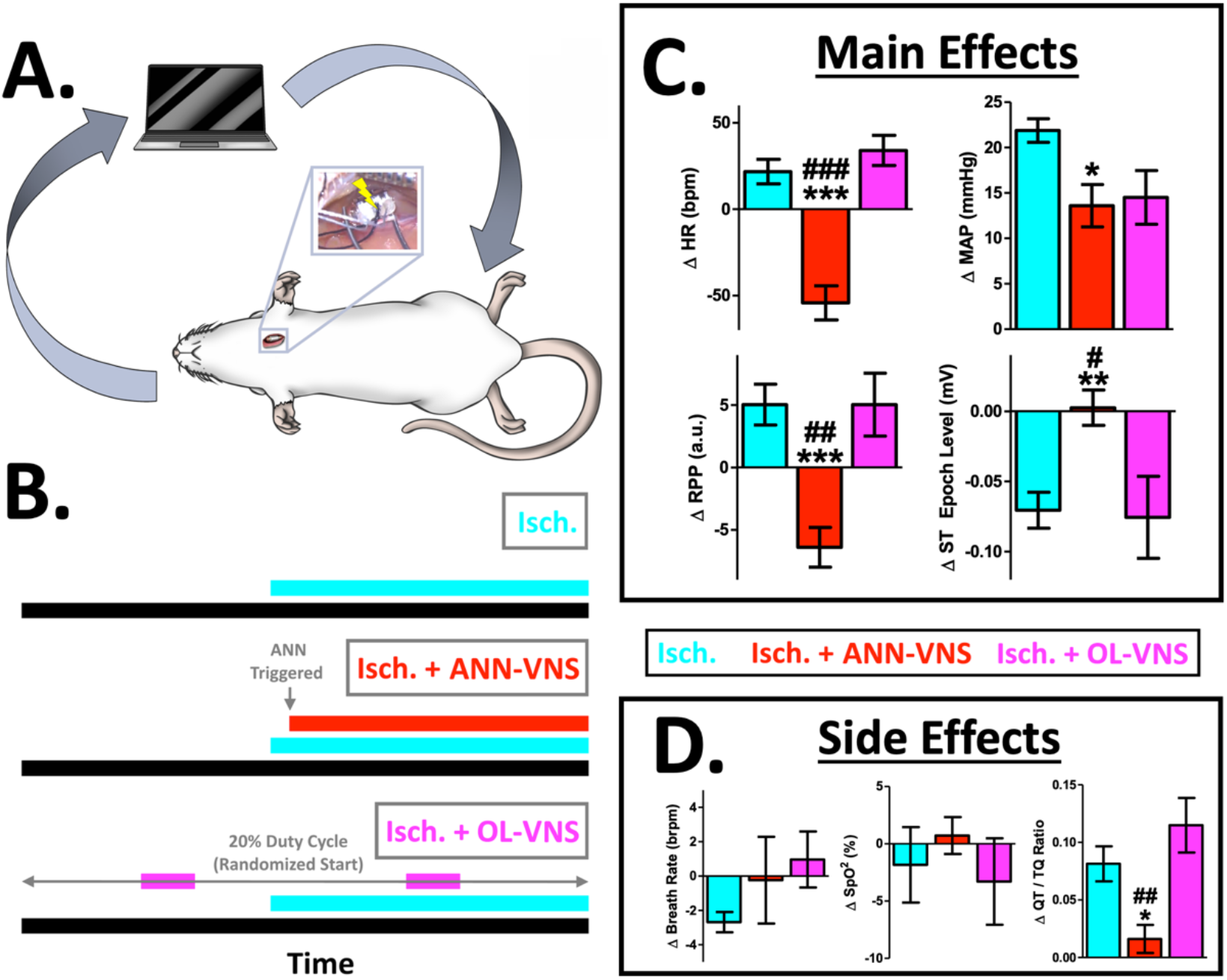
ANN Controlled Vagus Nerve Stimulation (ANN-VNS) Reverses Several Pathophysiological Correlates of Myocardial Ischemia Without Significant Side Effects. **A.** The ANN was next used on-line *in vivo* for rapid detection of spontaneous myocardial ischemia and control of vagus nerve stimulation (ANN-VNS; inset: left cervical vagus nerve and VNS cuff during dissection). **B.** We assessed biomarkers of cardiovascular stress and myocardial ischemia during either D+NE ischemia alone (cyan, Isch.), D+NE ischemia &closed-loop ANN-VNS (red, Isch. + ANN-VNS), and D+NE ischemia &open-loop VNS (magenta, Isch. + OL-VNS). Only closed-loop ANN controlled VNS (red, Isch. + ANN-VNS) reversed several biomarkers of myocardial ischemia, including heart rate, ST epoch level (electrophysiological correlate of subendocardial ischemia), rate-pressure product (RPP, index of myocardial oxygen consumption), and mean arterial pressure (MAP, correlate of afterload). Open-loop VNS (magenta, Isch. + OL-VNS) failed to reverse correlates of myocardial ischemia, and was essentially no different from myocardial ischemia alone (cyan, Isch.) (different from Isch. at: p<0.001 = ***, p<0.01 = **, or p<0.05 = *; different from Isch. + OL-VNS at: p<0.001 = ###, p<0.01 = ##, or p<0.05 = #). **D.** There were no significant differences in breath rate or blood oxygen saturation across groups (SpO^2^ = blood oxygen saturation). Importantly, only closed-loop ANN-VNS significantly mitigated a side effect related to arrythmia probability (QT / TQ ratio, averaged across ECG cycles). These results demonstrate the ability of ANNs to supplement biological neural networks and facilitate the reversal of spontaneous myocardial ischemia *in vivo* using a bioelectronic medicine. Data presented are mean ± SEM.

In real-time and *in vivo*, the ANN detected spontaneous D+NE induced myocardial ischemia with high overall accuracy (∼94%, Supplemental Fig. S5A; average decoder outputs: Supplemental Fig. S5B), similar to offline performance (∼92%). ANN-VNS reversed pathological changes in heart rate, MAP, RPP, and ST epoch level (Fig. 5C, Isch. + ANN-VNS, red), compared to D+NE ischemia alone (Fig. 5C, Isch., cyan; Heart Rate: F[2,20] = 29.6, p < 0.001; MAP: F[2,20] = 5, p < 0.05; RPP: F[2,20] = 14.2, p < 0.001; ST Epoch Level: F[2,20] = 7.6, p < 0.01; full 13-element feature vector shown in Supplemental Fig. S6A; experimental schematic: 4B). Open-loop VNS failed to reverse any major correlates of D+NE induced myocardial ischemia pathophysiology (Fig. 5C, Isch. + OL-VNS, magenta; open-loop VNS = 20% duty cycle, with a balanced amount of VNS compared to the closed-loop ANN-VNS paradigm, and parameters similar to previous human open-loop VNS studies: Anand et al., 2020; Table 1: Radcliffe et al., 2020; full 13-element feature vector shown in Supplemental Fig. S6B; experimental schematic: Fig. 5B, magenta). There were no significant side effects across groups related to breath rate or blood oxygen saturation (Fig. 5D; Breath Rate: F[2,20] = 0.9, p = 0.41; SpO^2^: F[2,20] = 0.4, p = 0.63). Importantly, only ANN-VNS significantly mitigated a side effect related to arrythmia probability (Fossa, 2017), and therefore enhanced myocardial electrical stability (Fig. 5D; QT / TQ ratio: F[2,20] = 9.3, p < 0.01). These findings demonstrate that pre-programmed open-loop VNS misses spontaneous myocardial ischemia that ANN-VNS is designed to respond to and correct. Overall, these results support the hypothesis that ANNs can supplement deficient biological neural networks in a number of ways: not only via detection, but also using bioelectronic control for correction of spontaneous cardiovascular pathophysiology.

Lastly, we performed a vagotomy caudal to the VNS site to examine the role of efferent vagal fiber activation. Caudal vagotomies blocked the major effects of ANN-VNS, indicating that the efferent fibers are critical for the therapeutic effects of ANN-VNS (Supplemental Fig. S6C, orange). Lastly, all 3 VNS groups received an equivalent amount of VNS (Supplemental Fig. S5C; F[2,14] = 1.3, p = 0.28). These additional findings show that both the vagal fibers engaged, and VNS timing (not necessarily VNS quantity), play critical roles in myocardial ischemia reversal.

## Detecting New Emerging Stress States Using ANN Autoencoders

Our next set of experiments addressed the need for decoding architectures to adapt as physiology changes. Over time, subjects can engage in new activities, and new forms of cardiovascular stress can emerge (Epel et al., 2018; Schwartz et al., 2018). A clinically deployed decoding architecture will fail if it is not capable of detecting new emerging physiological states. We assessed techniques potentially capable of detecting new, unknown, and emerging stress states (emerging state / outlier detection review: Park, 2019). To model new unknown emerging stress, we used feature data recorded during a higher magnitude of cardiovascular stress and myocardial ischemia (i.e., at a higher dose level; emerging stress states: H-D, H-NE, and H-D+NE). A subset of the detection techniques used an ANN approach (i.e., autoencoders).

LSTM autoencoders (LSTM-AE) detected new emerging stress states with a sensitivity of ∼99%, even though the network was not exposed to these states during training (Fig. 6A; F[2,27] = 26, p < 0.001; reconstruction loss distributions for known and unknown stress data: Supplemental Fig. S7; no significant differences for ‘known state’, i.e., D, NE, and D+NE, sensitivity across the 3 techniques, F[2,27] = 2.1, p = 0.13). Using sparse autoencoders removed the ability to assess long term dependencies in the data, significantly decreasing emerging stress state detection performance (i.e., no LSTM components, a Sparse-AE; Fig. 6A). The ANN enabled LSTM-AE also outperformed the widely used isolation forest technique (Fig. 6A). These results further demonstrate how biological neural networks can be supplemented with ANNs, and suggest that ANNs can also potentially adapt to new emerging physiological changes.

**Figure 6.**
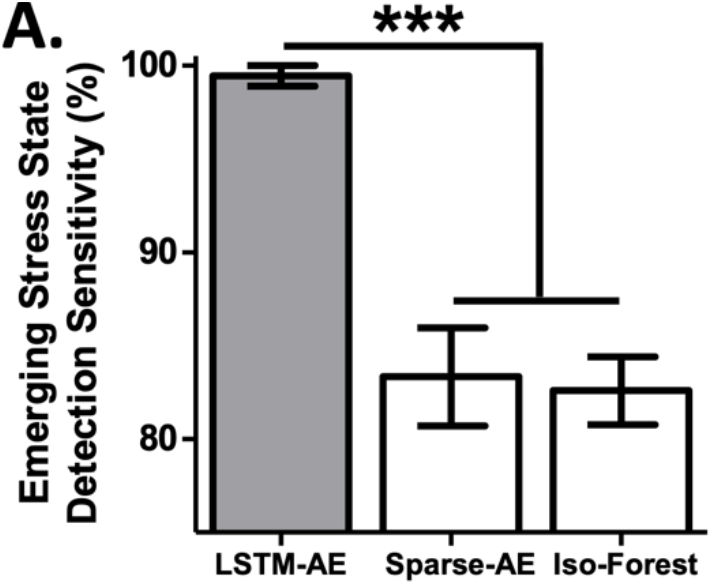
Detecting New Emerging Cardiovascular Stress States. **A.** We implemented techniques for detecting new emerging stress states (new emerging stress states = high dose versions of D, NE, and D+NE). LSTM autoencoders (LSTM-AE) significantly outperformed the sparse autoencoder (Sparse-AE) and isolation forest (Iso- Forest) approaches (*** = different at p<0.001). These results support the hypothesis that ANNs can also be used to detect new emerging stress states, as physiology evolves over time. Data presented are mean ± SEM.

### Enabling Interpretable and Adaptive AI: Visualizing Emerging Stress States Within the ‘Cardiovascular Latent Space’ and Unsupervised Dissociation of Different Emerging Stress Types

AI enabled medicines can suffer from a lack of interpretability – where either data or algorithm decisions cannot be readily understood. AI enabled medicines must be easily interpretable for widespread adoption (Vellido, 2019; Tonekaboni et al., 2019; Tjoa, E., &Guan, 2019). Visualizations are one solution for creating interpretable representations of both high-dimensional data and complex algorithm decisions.

We next created an interpretable visualization of all known and new emerging stress states (Fig. 7A; using the LSTM-AE hidden layer, and the dimensionality reduction technique uniform manifold approximation and projection, or UMAP; McInnes &Healy, 2018). This architecture approach has recently achieved state-of-the-art performance converting complex high dimensional data into interpretable representations (McConville et al., 2019). Across all stress states, the uninterpretable high dimensional LSTM-AE hidden layer (256 dimensions) was transformed to an interpretable 2-dimensional representation (Supplemental Video 1, Fig. 7B, &Supplemental Fig. S8). In this ‘cardiovascular latent space’, known and new emerging states formed clear clusters (Fig. 7B). Furthermore, known and new emerging stress states generally occupied separate regions of the ‘cardiovascular latent space’ at ∼85% accuracy, performing well above chance levels (t[18] = 7.1, p < 0.001; data from all 10 folds: Supplemental Fig. S8; performance = ability to separate the known and new emerging stress state clusters using a linear boundary). This interpretable visual information showcases the ability of ANNs to help meet clinical needs and create meaningful representations of complex high-dimensional cardiovascular data, even though the architecture has never been exposed to the new emerging stress state data.

**Figure 7.**
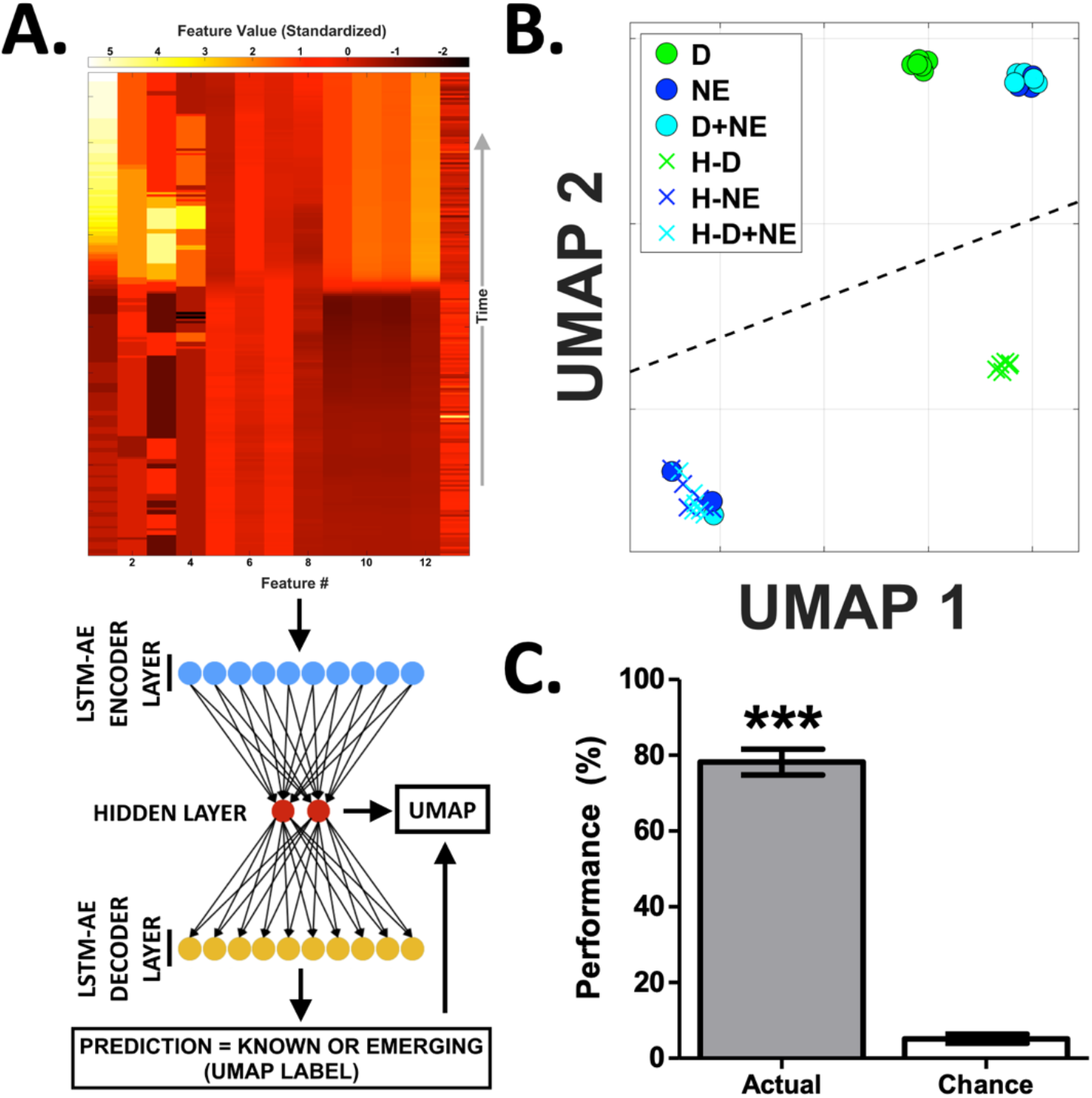
Leveraging the ‘Cardiovascular Latent Space’ for Unsupervised Identification of New Emerging Cardiovascular Stress States. **A.** Schematic of the emerging state detection architecture. For a given stress state observation, the feature matrix (top) is passed into the LSTM-AE (consisting of the encoder, hidden layer, and decoder components; LSTM-AE cartoon not to scale). Using the reconstruction loss, the LSTM-AE then predicts whether the given observation is a known or new emerging stress state. This prediction is then used as a label for UMAP dimensionality reduction and subsequent visualization of the LSTM-AE’s processes (see Supplemental Video 1 for a representative movie of these processes). **B.** The hidden layer of the LSTM-AE is an uninterpretable 256-dimensional vector. We next generated an interpretable version of all known and unknown stress states using a combination of the LSTM-AE hidden layer and the dimensionality reduction technique UMAP (uninterpretable input = 256 dimensions; interpretable output = 2 dimensions). This ‘cardiovascular latent space’ interestingly contained clustering of known (circles) and unknown emerging stress states (X’s), indicating that the ‘cardiovascular latent space’ may also be useful for identifying different types of new emerging stress states (plotted data is representative of overall performance; black dashed line: linear decision boundary, calculated using a SVM). **C.** We next combined the ‘cardiovascular latent space’ with unsupervised clustering to potentially identify different types of new emerging stress states (via hierarchical density based spatial clustering of applications with noise, or HDBSCAN). This fully unsupervised method achieved 78% performance when challenged to autonomously detect emerging stress states, performing well above chance performance levels (*** = different from chance at p<0.001). Data presented are mean ± SEM. Overall, these findings show that an ANN can help enable an interpretable and unsupervised emerging state detection architecture, relevant for a detection system that can adapt as physiology changes.

Emerging state detection architectures should also be able to autonomously identify different types of emerging states, if multiple types exist. Unfortunately, this is exceedingly challenging, as the architecture cannot be exposed to one or multiple types of new emerging stress states during training. To address this challenge, our final analyses leveraged the LSTM-AE enabled ‘cardiovascular latent space’ combined with unsupervised clustering (unsupervised clustering method: hierarchical density-based spatial clustering of applications with noise or HDBSCAN; Campello et al., 2013). This fully unsupervised architecture achieved 78% performance when challenged to autonomously identify different types of emerging stress states, performing well above chance levels (Fig. 7C; i.e., unsupervised dissociation of varying combinations of H-D, H-NE, and H-D+NE; t[18] = 20.1, p < 0.001; performance metric = V-measure * 100%, Rosenberg &Hirschberg, 2007; V-measure is a well-studied metric for quantify clustering and detection capability; Supplemental Fig. S9: separate completeness, homogeneity, and performance values for all 7 emerging stress state scenarios). The architecture achieved this performance, in spite of unsupervised operation and significant out of sample generalization to multiple types of new emerging stress states. Overall, these results show that ANNs can further enable an interpretable unsupervised emerging state detection architecture, relevant for adapting to physiological changes over time.

## Discussion

“It is hard to understand the biological strategy and hence development of a system providing the wild animal with hundreds of fibers exclusively designed for signaling unlikely coronary emergencies”

-Alberto Malliani (Malliani, 1986)

In this study, we demonstrate several ways ANNs can supplement deficient biological neural networks. ANNs effectively decoded cardiovascular states with high accuracy, even though biomarkers exhibited significant variability and state overlap, similar to human myocardial ischemia. Beyond detection alone, an ANN enabled bioelectronic medicine reversed myocardial ischemia by reactively triggering VNS to reduce correlates of chronotropy, afterload, and myocardial oxygen demand. Preprogrammed open-loop VNS or ANN-VNS without efferent vagal fibers intact both failed to reverse myocardial ischemia, demonstrating the importance of VNS timing and vagal fibers engaged. Lastly, ANNs enabled clinically relevant interpretable visualizations and adaptive detection of emerging cardiovascular stress. This study demonstrates for the first time that ANNs can supplement deficient cardiovascular biological neural networks via an artificially intelligent bioelectronic medicine system.

### Supplementing Deficient Biological Neural Networks with Artificial Neural Networks

It is exceedingly problematic that the leading cause of mortality world-wide - cardiovascular disease and myocardial ischemia - largely develops without conscious awareness.∼75% of myocardial ischemia events are asymptomatic and therefore subperceptual, known as ‘silent myocardial ischemia’ (Gutterman, 2009; Deedwania &Nelson, 1990; Rozanski &Berman, 1987; Cecchi et al., 1983). Furthermore, up to ∼50% of myocardial infarctions (i.e., ‘heart attacks’) are also asymptomatic and happen without any sensation (Soliman, 2019). These significant deficits in biological neural networks likely come from several sources.

Firstly, deficient detection of myocardial ischemia may be due to evolutionary constraints. Several human-related factors that contribute to cardiovascular disease are relatively new from an evolutionary perspective, including consuming high-fat foods, smoking, or a sedentary lifestyle (Ding &Kullo, 2009). Therefore, there may have been insufficient time to develop an effective cardiovascular pathophysiology detection system in humans, via evolutionary modifications to neural systems or other mechanisms (Kember et al., 2013; Ding &Kullo, 2009; Malliani, 1986).

Secondly, ischemia itself and other diseases contribute to deficient detection of myocardial ischemia. Symptomatic ischemia, known as angina, only comprises ∼25% of all ischemic events and is often misdiagnosed as off-target musculoskeletal pain, making it difficult to diagnose accurately (Gutterman, 2009; Swap &Nagurney, 2005). Even when angina occurs, subsequent ischemic events can be silenced and become asymptomatic, via desensitization of afferent signaling (known as ‘neural stunning’: Gutterman, 2009; Pomblum et al., 2010). Lastly, cardiovascular disease can accompany other disorders such as diabetes. Diabetic autonomic neuropathy further impairs myocardial ischemia signaling, degrading neural sensing systems innervating the heart (Tabibiazar &Edelman, 2003; Pop-Busui, 2010).

Regardless of the mechanism, biological neural networks are largely incapable of reliably detecting myocardial ischemia. In this study, we address this deficiency of biological neural networks using ANNs. Future approaches that supplement biological neural networks using ANNs hold significant promise for mitigating numerous shortcomings of physiological systems.

Biological neural networks and ANNs have had a long history together. These interactions range from the early days of parallel distributed processing to recent ANN architectures that mimic mammalian neural systems (reviews: Hassabis et al., 2017; Marblestone et al., 2016). Aside from controlling a therapeutic device during disease, ANNs can now also assist healthy humans (e.g., medical diagnoses, self-driving cars, military applications, and more: Wilson &Daugherty, 2018; Jarrahi, 2018). These findings further highlight several areas of opportunity for ANNs to enhance human function, during either disease or even healthy states.

### Reversing Spontaneous Myocardial Ischemia Using A Responsive Closed-loop Bioelectronic Medicine

High rates of ‘silent myocardial ischemia’ lead to increases in myocardial injury, myocardial infarction, and sudden death (Conti et al., 2012; Gutterman, 2009; Lotze et al., 1999; Deedwania &Carbajal, 1990). Treating silent or symptomatic myocardial ischemia reduces rates of myocardial injury, myocardial infarction, and death (Braun et al., 2018; Conti et al., 2012; Gutterman, 2009; Cohn, 1998). Treating myocardial ischemia using a pharmacological medicine can promote vasodilation and / or reestablish an appropriate myocardial oxygen supply-demand ratio (Balla et al., 2018; Cohn, 1998). VNS mimics these desired effects via cholinergic modulation, decreasing intracellular calcium, presynaptic inhibition of norepinephrine release, coronary vasodilation, and other mechanisms (Capilupi et al., 2020; Ardell et al., 2015; Levy &Schwartz, 1994). Bioelectronic control of these physiological cascades motivated the use of VNS in this study.

Bioelectronic medicines are beginning to address several shortcomings of pharmacological medicines (Ganzer &Sharma, 2019; Vitale &Litt, 2018; Birmingham et al., 2014). Although pharmacological medicines can be effective, they do not target specific tissues leading to side-effects. Bioelectronic medicines can address this limitation, via stimulating specific nerves, and therefore targeting specific tissues, for a localized effect. Furthermore, several disease episodes are spontaneous and may only occur for several minutes a day. Bioelectronic medicines can be dynamically switched on and off as needed, unlike pharmacological medicines that are active for several hours a day. Importantly, bioelectronic medicines can provide on-demand benefit via closed-loop activation. This on-demand attribute of bioelectronic medicine can reduce desensitization of target receptors, further mitigate side effects, and ultimately improve therapeutic efficacy. Overall, bioelectronic medicines mitigate several shortcomings of pharmacological medicines, providing spatial and temporal specificity to improve therapeutic outcomes and reduce off target effects.

Our findings extend previous studies that apply vagal modulation during myocardial ischemia (Machada et al., 2020; Nuntaphum et al., 2018; Del Rio et al., 2008; Vanoli et al., 1991; Buck et al., 1981; Meyers et al., 1974), and specifically highlight the importance of responsive closed-loop VNS control (Fig. 5 and Supplemental Fig. S6). Notably, only responsive closed-loop VNS (with efferent vagal fibers intact) reversed major correlates of myocardial ischemia (Fig. 5 and Supplemental Fig. S6). Efferent cervical vagal fibers innervate both the atria and ventricles (Capilupi et al., 2020; Levy &Schwartz, 1994). Acetylcholine release from efferent vagal fibers can mitigate elevated chronotropy, inotropy, afterload, and myocardial oxygen consumption seen during myocardial ischemia (Capilupi et al., 2020; Levy &Schwartz, 1994; Ardell et al., 2015; Nuntaphum et al., 2018; Del Rio et al., 2008; Vanoli et al., 1991; Buck et al., 1981; Meyers et al., 1974). Our results demonstrate that closed-loop intact VNS decreases overall myocardial work, important for preventing cell death and injury during myocardial ischemia.

During myocardial ischemia alone, we observed depression of ECG segments during both systole and diastole (Fig. 3). These ECG epochs depress during the initial stages of myocardial ischemia, indicative of ischemic currents (Klabunde, 2017; Janse, 2007; Cinca et al., 1980; Kleber et al., 1978). Closed-loop intact VNS reduced chronotropy, afterload, myocardial oxygen consumption, and other factors leading to a full reversal of ST epoch depression. This result importantly demonstrates the complete reversal of subendocardial ischemia (Klabunde, 2017) during closed-loop intact VNS (Fig. 5). These findings support the hypothesis that closed-loop intact VNS suppresses these ischemic currents, via a responsive increase in parasympathetic drive restoring myocardial oxygen balance.

Lastly, the cardiovascular effects of VNS required precise timing and delivery of stimulation during spontaneous ischemic episodes. Open-loop VNS was not programmed to respond during spontaneous myocardial ischemia, and thus generally failed to affect biomarkers of myocardial ischemia (Fig. 5 and Supplemental Fig. S6). Therefore, open-loop VNS may simply miss random myocardial ischemia events. We used an open-loop VNS paradigm representative of human cardiovascular studies, near the upper limit of clinically tolerable VNS levels (20% duty cycle at 2-2.5 mA; Anand et al., 2020; Table 1: Radcliffe et al., 2020). From a translational perspective, the closed-loop VNS paradigm used here should deliver significantly less VNS compared to open-loop VNS over time. For example, several clinical studies indicate that myocardial ischemia can occur for several minutes and up to ∼1 hour per day (Pepine et al., 1994; Trimarco et al., 1990; Hinderliter et al., 1991). Therefore, to responsively mitigate myocardial ischemia, closed-loop VNS may only be needed for ∼1 hour a day or less. Over a 24-hour period, closed-loop VNS should also deliver ∼1-2 orders of magnitude less VNS compared to open-loop VNS. The total charge delivery of VNS is important for future safety studies aiming to treat spontaneous myocardial ischemia with VNS. These results motivate future studies to optimize the total amount of stimulation delivered using responsive bioelectronic medicines, keeping in mind the desired safety and efficacy.

### AI Enabled Medicines: Opportunities and Challenges

State-of-the-art machine learning methods provide powerful capabilities for pattern recognition that in many cases exceed the abilities of expert humans. The financial industry was an early adopter of neural network models for forecasting stock market index, energy demand, and real estate prices, prompted initially by a need to model nonlinear multivariate datasets (Huang et al., 2007; Wang et al., 2018). In medicine, the role of AI has been increasing steadily (Miller &Brown, 2018), especially in the field of Radiology where AI-enabled systems are used not only for detection and interpretation of images, but also scheduling and triage, clinical decision support systems, and several other critical steps of the Radiology workflow (Choy et al., 2018).

The uptake of AI-based solutions is driven by their capacity to ingest and comprehend vast quantities of data, permitting a more comprehensive assessment of a patient’s condition. Included in this is the ability to detect dynamic features that are not apparent in the typical snapshot evaluations that are performed in the clinic (Romiti et al., 2020; e.g., blood pressure and heart rate at a single point in time). We leverage these capabilities of AI systems to dynamically detect and correct pathological cardiovascular events *in vivo* (Fig. 5 and Supplemental Fig. S6), similar to previous studies using responsive therapies for cardiovascular treatment (Kawada, T., &Sugimachi, 2009; Gotoh et al., 2005; Sugimachi, M., &Sunagawa, 2009; Sato et al., 2002).

Despite the clear benefits of AI-enabled technology solutions, trustworthiness is a major barrier to the adoption of AI-based diagnostics, and especially intervention. Some patients and physicians may be reluctant to allow a computer to make healthcare decisions. A recent survey of radiologists, information technology specialists, and industry representatives found that only 25% of the 123 people surveyed expressed confidence in results obtained by AI systems used in Radiology, and the vast majority (∼91%) emphasized the need to validate the algorithms used in these systems (Jungmann et al., 2020). Strategies for building trust include the creation of ‘Explainable AI’ that provides greater transparency and traceability, especially for systems that rely on deep learning architectures that are particularly opaque (Holzinger et al., 2019; Tjoa, E., and Guan, 2020). Improved methods for data and model visualization may facilitate interpretability and explainability in medical AI systems (Vellido, 2019, 2012, &2011; Liu et al., 2017), and were leveraged in the current study (via autoencoders, dimensionality reduction, and unsupervised clustering; Fig. 6 and 7). Importantly, building trust will likely be achieved gradually through an evolution of clinical trials that demonstrate with hard evidence the benefits of AI-based approaches in improving patient care.

## Supplemental Figures

**Supplementary Figure S1.**
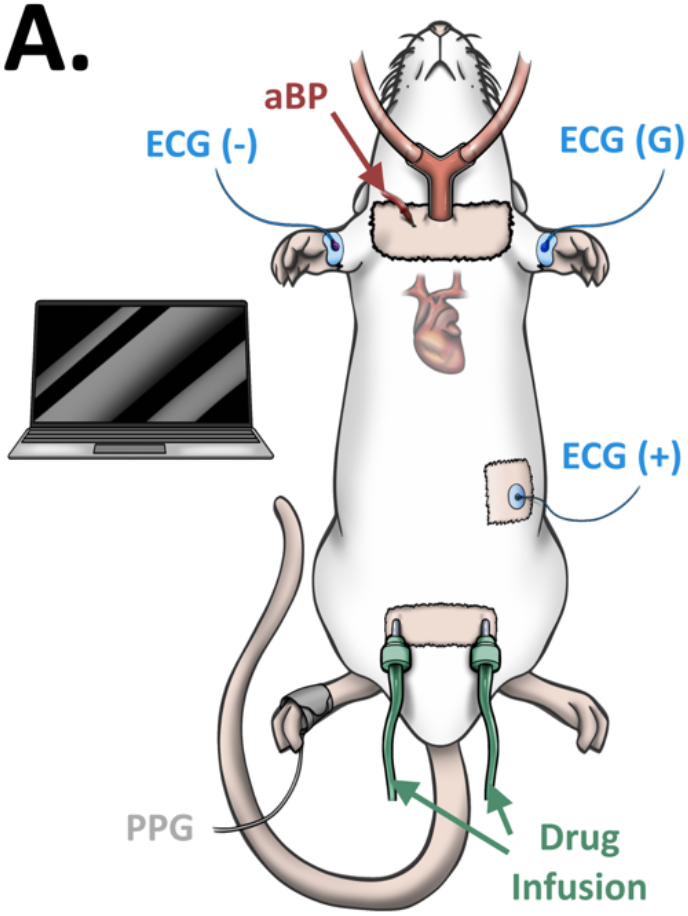
Cartoon Schematic of Experiment (related to Fig. 1, 2, &3). **A.** Cartoon schematic of the *in vivo* experiment and interfaces. All experiments were performed in isoflurane anesthetized rats (using tracheotomy, light red tube). We recorded arterial blood pressure from within the right carotid artery (aBP, red), a lead II electrocardiogram (ECG, blue patches; negative, positive, and ground electrodes noted), and a photoplethysmogram (right foot, black patch) during injections of cardiovascular stress and myocardial ischemia inducing agents into the femoral veins (catheters, green). All modules were synchronized and controlled electronically (laptop computer).

**Supplementary Figure S2.**
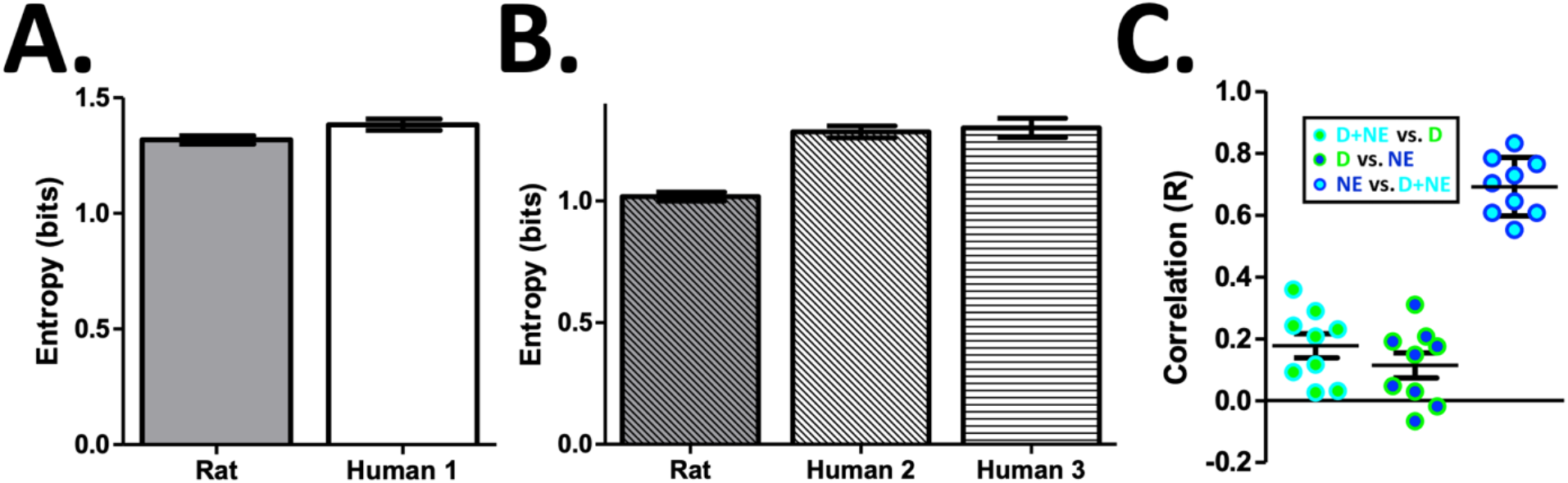
Cardiovascular Feature Data Exhibit Variability and State Overlap, Similar to Human Stress and Myocardial Ischemia (related to Fig. 3). A &B. Cardiovascular feature data recorded from the rat (‘Rat’) demonstrated similar levels of variability and entropy compared to cardiovascular feature data recorded from human subjects in either the intensive care unit (**A**; including all 13 features, ‘Human 1’) or from human subjects during ambulatory myocardial ischemia (**B**; including only ECG features, #1 - #8, ‘Human 2’ and ‘Human 3’). **C.** Cardiovascular feature data from the rat also exhibited significant stress state overlap, specifically between the NE and D+NE states. These results support the hypothesis that cardiovascular stress and myocardial ischemia induced by D, NE, and D+NE injections induce variability and state overlap in the cardiovascular data, similar to human cardiovascular stress states. Data presented are mean ± SEM.

**Supplementary Figure S3.**
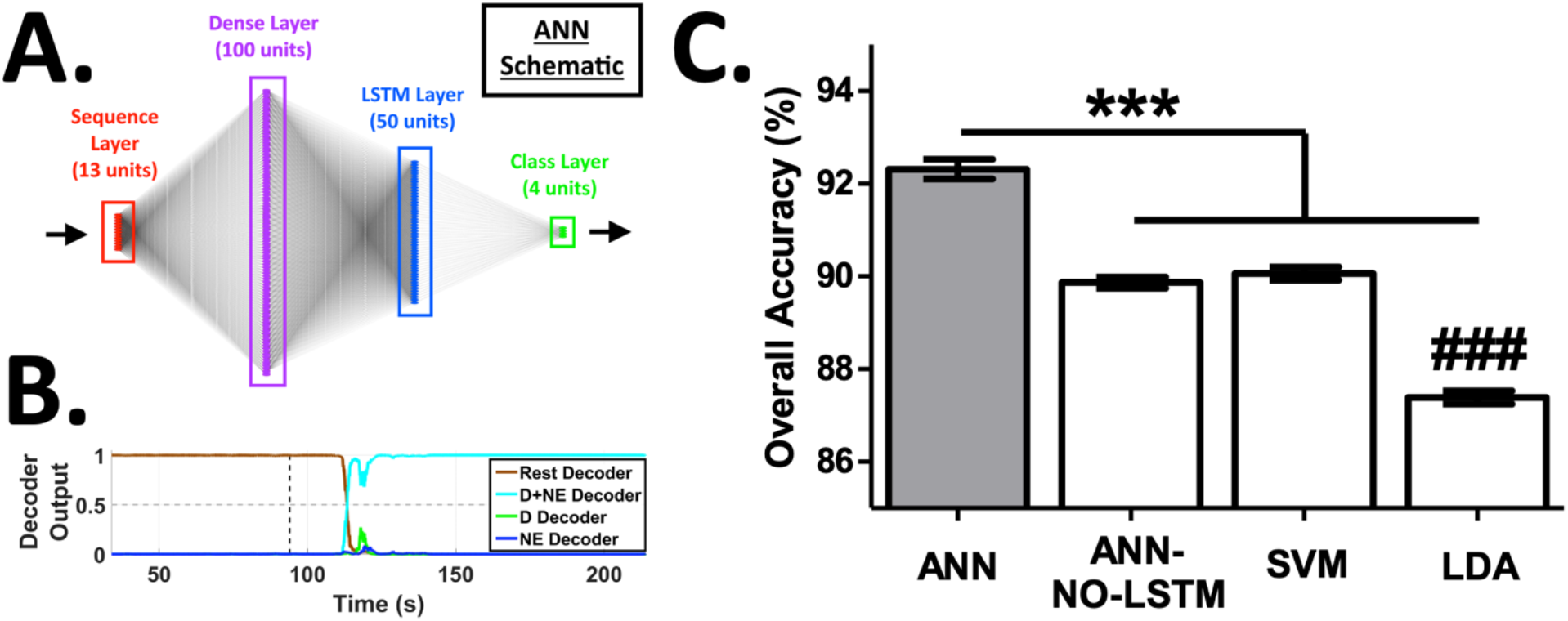
Artificial Neural Network (ANN) Architecture, ANN Decoder Outputs, and Superior Performance Compared to Other Classifiers (related to Fig. 4). **A.** Cartoon schematic of the 4-layer ANN architecture (layers not to scale; red: sequence input layer, 13 units; purple: dense layer, 100 units; blue: LSTM layer, 50 units; green: class output layer, 4 units). Please see the *Decoding Myocardial Demand Ischemia Using an Artificial Neural Network (ANN)* section of the methods for more details on the ANN. **B.** A given recording begins with a 90 s period of rest (i.e., no drug injected) followed by a 120 s period of the given injected agent. Feature creation and decoding began at 34 seconds to allow for the recording of sufficient baseline activity. Example ANN decoder outputs across the 4 classes during an injection of D+NE (a respective decoder output ranges from zero [low confidence in the respective class] to 1 [high confidence in the respective class]; gray dashed line = decoder significance threshold; black dashed line = injection start). **C.** The ANN outperformed an artificial neural network without an LSTM layer (ANN-NO-LSTM), a support vector machine (SVM), and a linear discriminant analysis (LDA) (*** different at p<0.001; ### different from ANN-NO-LSTM or SVM at p<0.001). These results demonstrate the superior performance of ANNs and the importance of leveraging time series dependencies for cardiovascular state decoding. Data presented are mean ± SEM.

**Supplementary Figure S4.**
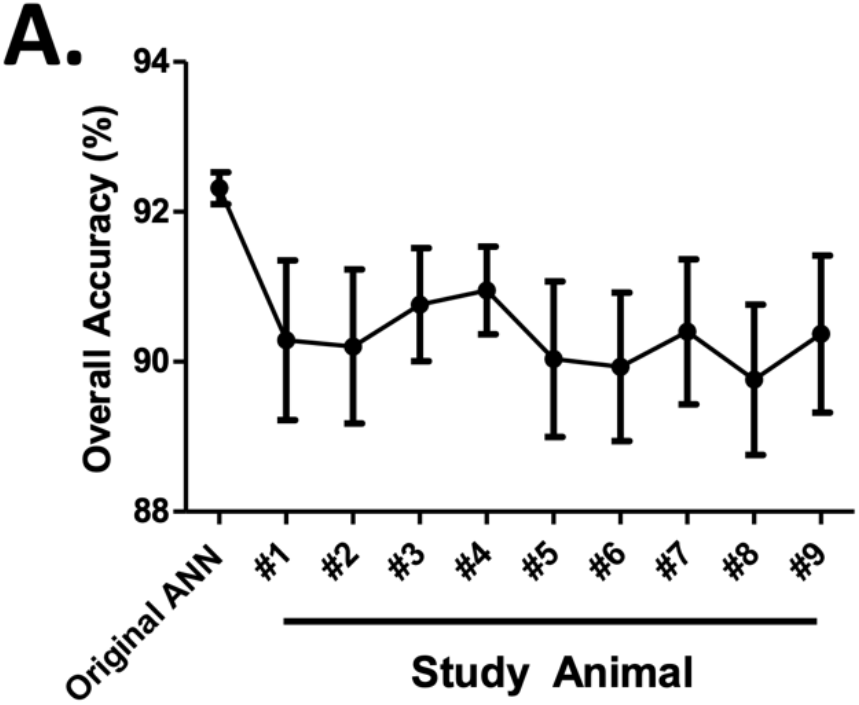
Fixed ANNs Demonstrate Significant Robustness and Generalization Out of Sample Across Time and Animals (related to Fig. 4). **A.** Fixed ANNs were created to assess model generalization well out of sample to animals across the entire study. The original ANN performance level is shown as a reference (left, ‘Original ANN’), where the model was trained on data from the whole study and therefore all animals. The remaining performance levels are shown for separate ‘fixed’ ANNs. A fixed ANN was first trained on the base data set plus the given animal’s data. The given fixed ANN was then challenged to predict on the remaining animals in the study (several weeks into the past or future), without any model updating. Although there was a decrease in accuracy and an increase in prediction variance, fixed ANNs still generalized well across time and even to other animals (chance level of prediction = ∼25%). These results indicate that ANNs are robust and can generalize well out of sample. Data presented are mean ± SEM.

**Supplementary Figure S5.**
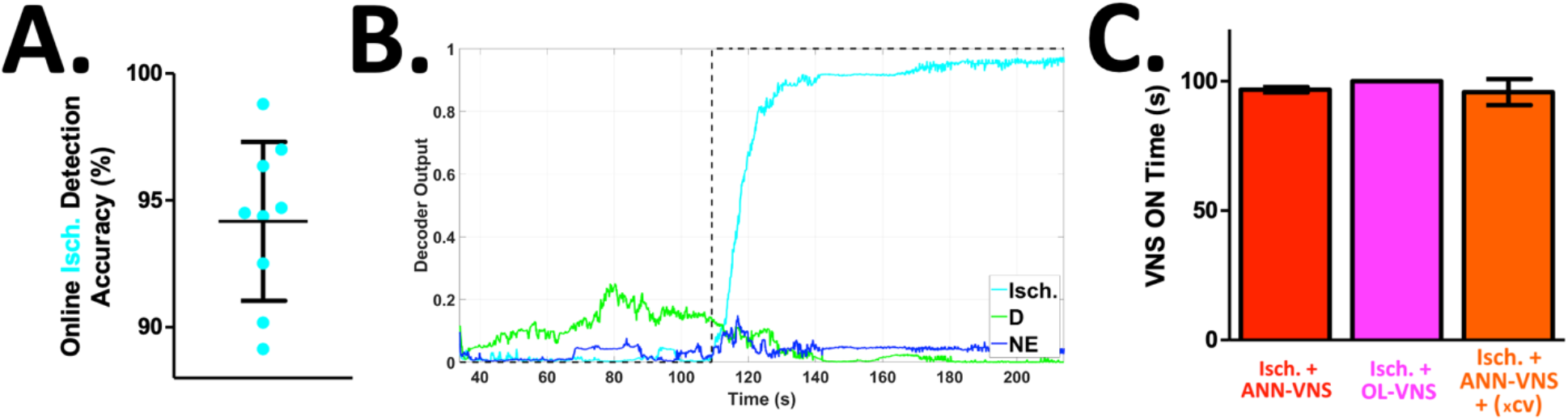
*In vivo* ANN Decoding Performance, Average ANN Decoder Outputs, and VNS Quantity Across Groups (related to Fig. 5). **A.** The ANN performed *in vivo* online decoding of the target ischemic state (i.e., a D+NE injection) with an overall accuracy of ∼94% (cyan points = overall accuracies from single animals). **B.** ANN decoder outputs for the 3 cardiovascular stress states averaged across all animals from the *in vivo* experiments (N = 9; black dashed line = labeled period for the given injection). **C.** All 3 VNS groups received the same quantity of VNS (red = Isch. + ANN-VNS; magenta = Isch. + OL-VNS; orange = Isch. + ANN-VNS + (xcv)). Data presented are mean ± SEM.

**Supplementary Figure S6.**
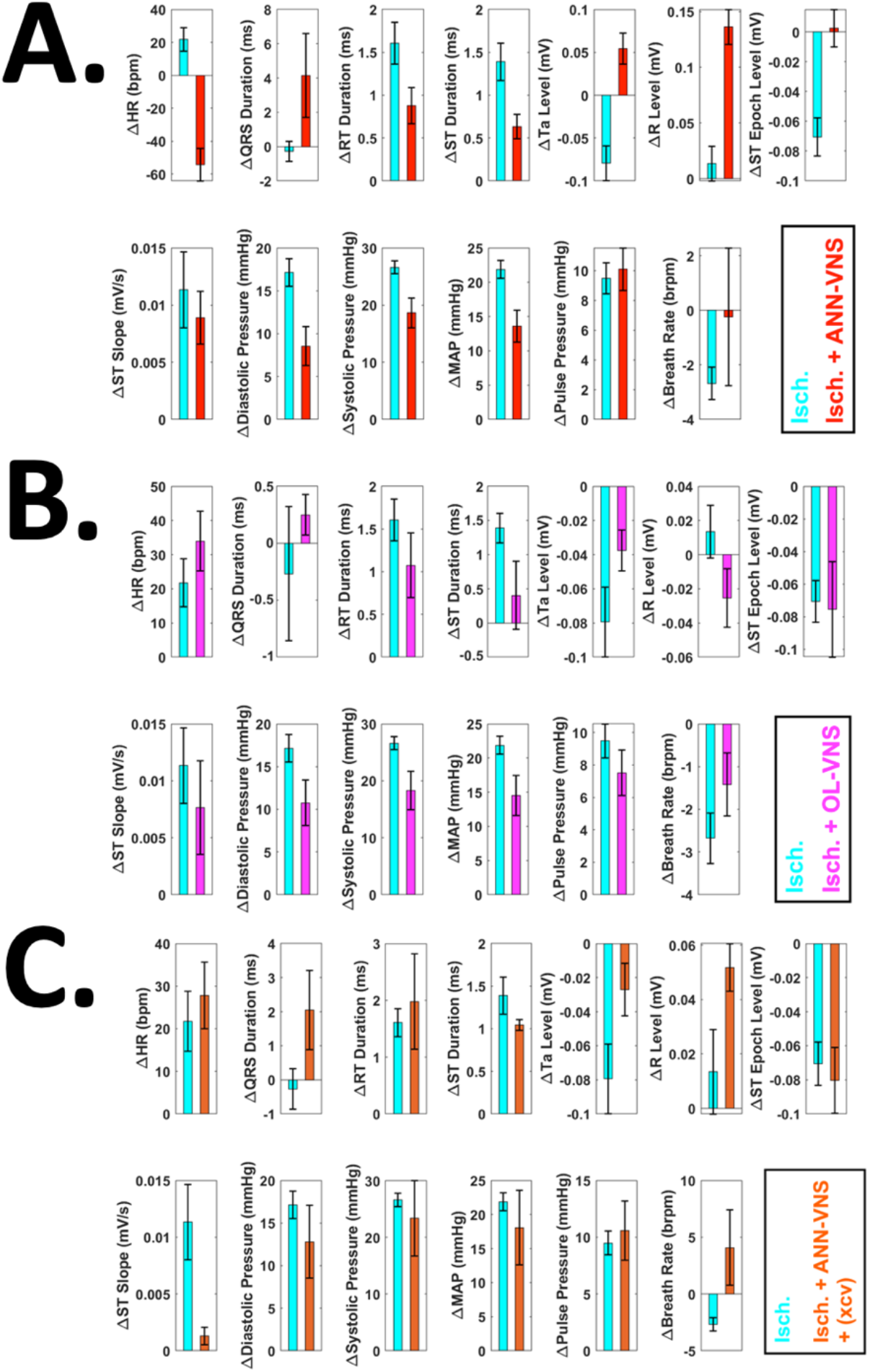
Effects Across All 13 Features Using Either Closed-loop VNS, Open-loop VNS, or Closed-loop VNS Following a Vagotomy Caudal to the VNS site (related to Fig. 5). **A.** All 13 features during either D+NE ischemia alone (cyan, Isch.) or D+NE ischemia &closed-loop ANN-VNS (red, Isch. + ANN-VNS). We performed 2 controls to appraise the mechanism of ANN-VNS. Both preprogrammed open-loop VNS (all features: **B**; magenta, Isch. + OL-VNS) and ANN controlled VNS following a vagotomy caudal to the VNS site (all features: **C**; orange, Isch. + ANN-VNS + (xcv)) essentially failed to significantly affect cardiovascular pathophysiology induced by ischemia alone (cyan, Isch.). The results highlight the importance of both closed-loop VNS and vagal fibers engaged for reversing myocardial ischemia pathophysiology. Data presented are mean ± SEM.

**Supplementary Figure S7.**
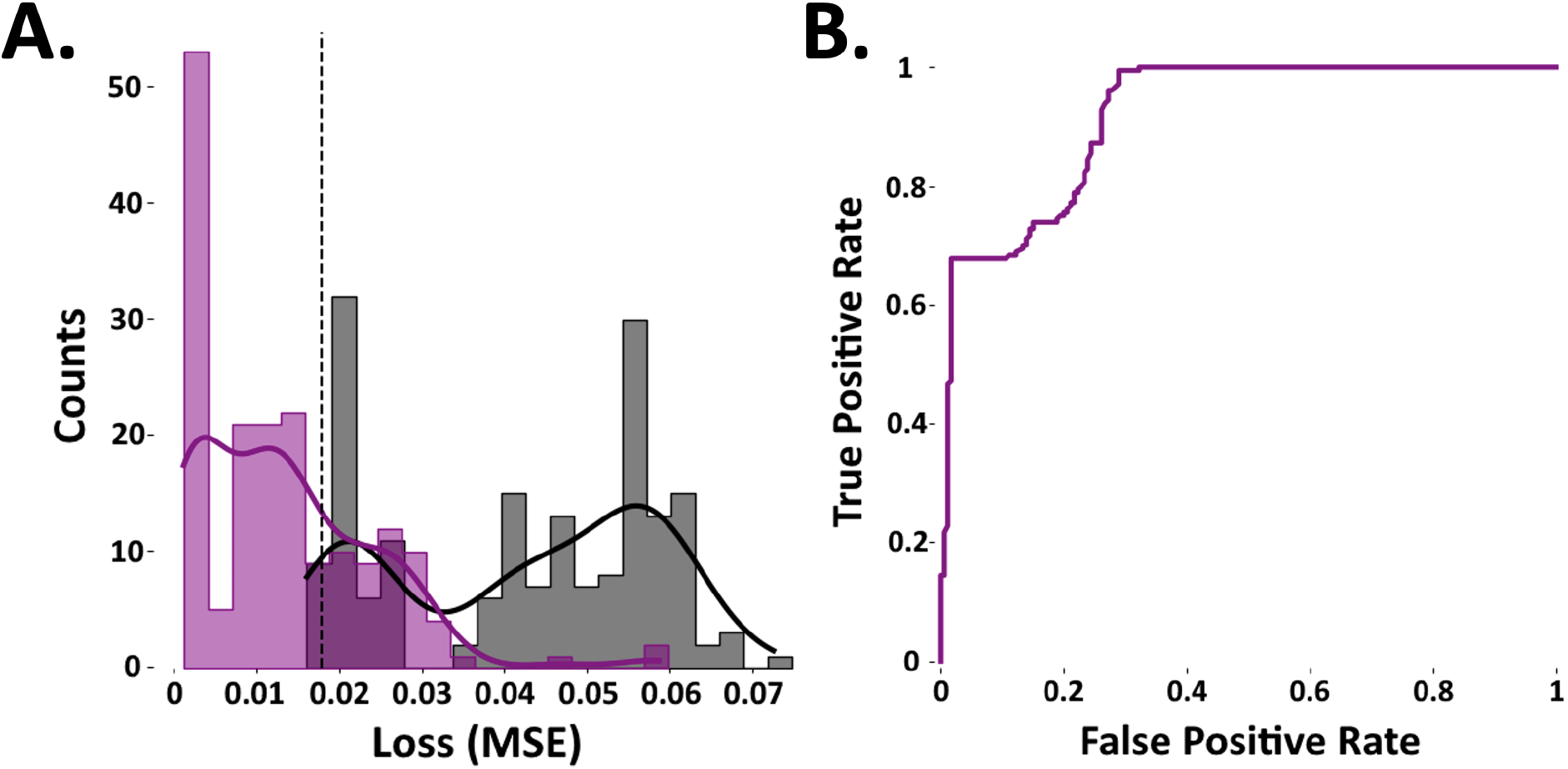
Emerging Cardiovascular Stress State Detection: Raw Mean Square Error (MSE) Reconstruction Distributions for The LSTM Autoencoder (LSTM-AE) (related to Fig. 6). Using the LSTM-AE, emerging stress states were detected using a simple threshold method related to the reconstruction loss (i.e., mean square error or MSE), similar to previous studies (Park et al., 2019). A high reconstruction loss is indicative of a new emerging state that has never been seen by the LSTM-AE, and a low reconstruction loss indicates that the state is known. The MSE loss distributions across all folds are shown for the LSTM-AE models for reconstruction of known stress states (**A**, purple; i.e., D, NE, and D+NE), or new emerging stress states (**A**, gray; i.e., H-D, H-NE, and H-D+NE). Colored curves (gaussian kernel fits) are shown on top of a given distribution (vertical dashed line = MSE threshold for determining known and new emerging stress states, optimized for accuracy). **B.** Receiver operating characteristic curve for the LSTM-AE technique, where true positive and false positive rates are plotted across a range of MSE thresholds.

**Supplementary Figure S8.**
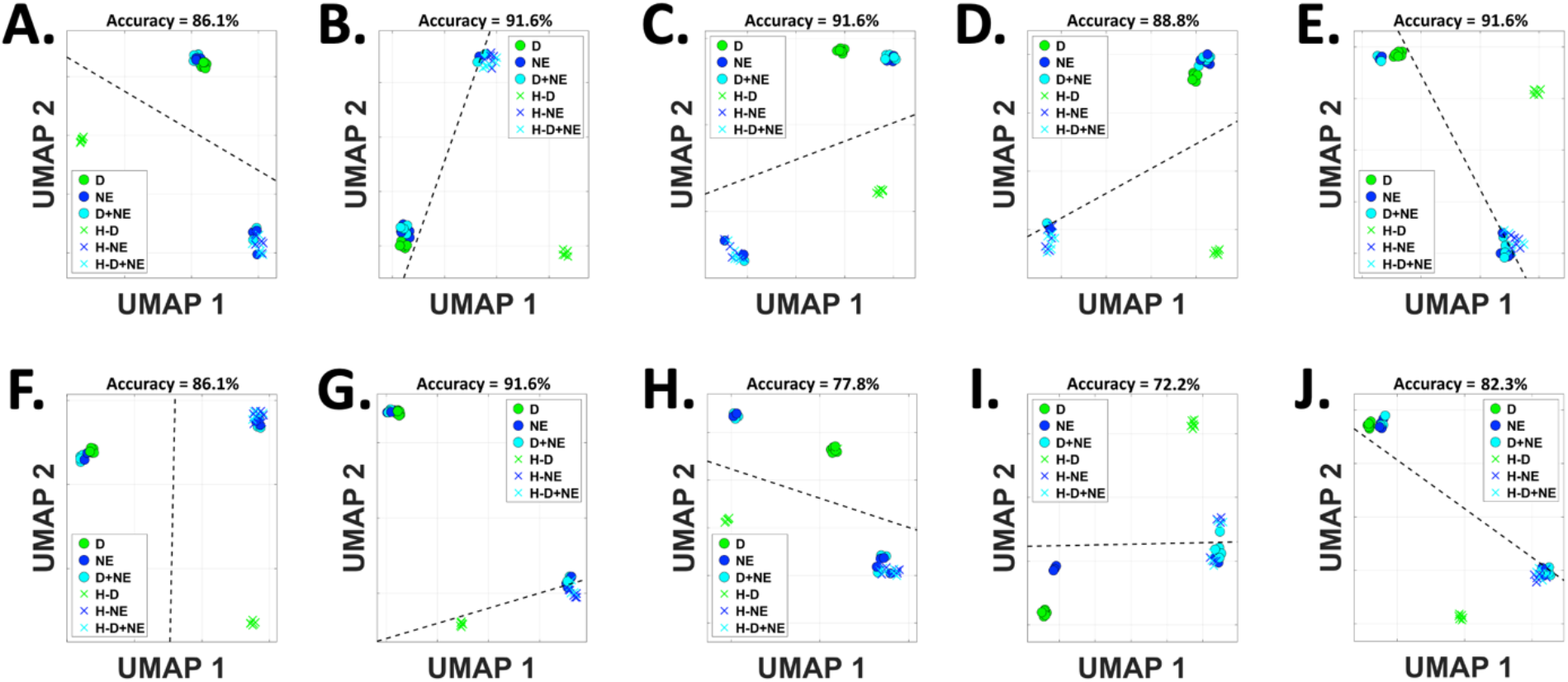
Known and Unknown Cardiovascular Stress States Within the ‘Cardiovascular Latent Space’: Raw Data from All Folds (related to Fig. 7B). We assessed 2-dimensional representations of all known and unknown stress states within the ‘cardiovascular latent space’, leveraging the emerging state identification architecture (architecture schematic: Fig. 7A; known stress states: D, NE, and D+NE; unknown stress states: H-D, H-NE, and H-D+NE). Known and unknown stress states generally occupied mutually exclusive regions of the ‘cardiovascular latent space’ across all folds at ∼85% accuracy (folds 1-10 = panels **A – J**, respectively; region boundary: black dashed line, determined using a linear support vector machine; separation accuracy shown above each plot). These results demonstrate the ability to robustly increase interpretability and accurately visualize known and new unknown emerging stress states.

**Supplementary Figure S9.**
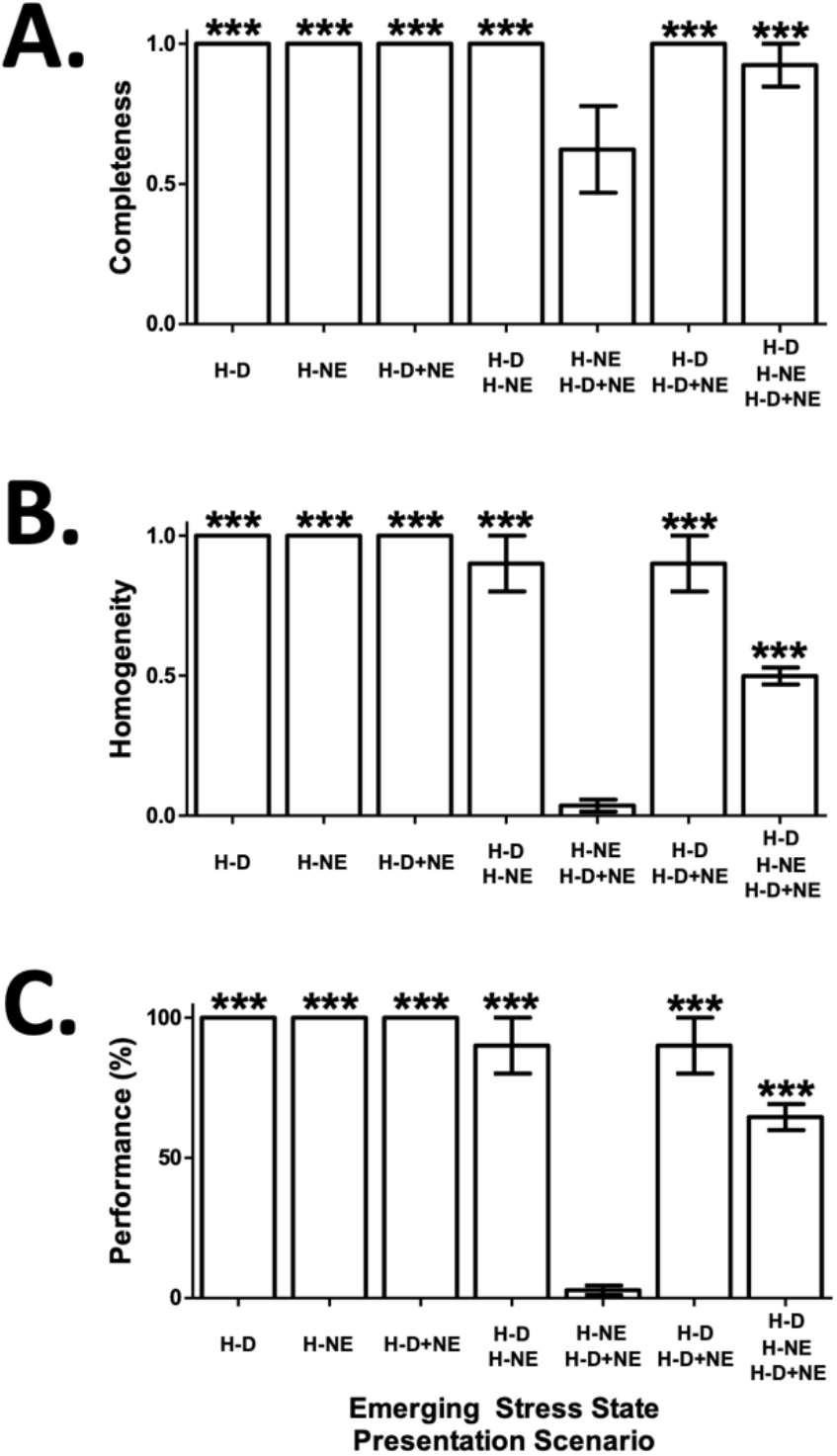
Emerging Cardiovascular Stress State Detection Performance Metrics Across All 7 Presentation Scenarios (related to Fig. 7C). We performed emerging stress state detection across the 7 emerging stress state presentation scenarios (x-axes), and present well studied correlates of unsupervised clustering performance (Rosenberg &Hirschberg, 2007) including completeness (**A**), homogeneity (**B**), and performance (V-measure * 100%, **C**). Across metrics, almost all presentation scenarios performed well above chance performance levels (*** = different from chance at p<0.001). Data presented are mean ± SEM.

## Acknowledgements

We would like to thank our development and management teams at Battelle Memorial Institute, Russ Kittel for his contributions to the manuscript graphics, and several collaborators who contributed to the preparation and review of the manuscript. Financial support for this study came from Battelle Memorial Institute.

## Author Contributions

P.D.G., S.R.R., B.T., W.W.M., D.J.W., and R.L.H. conceived and designed the experiments. P.D.G., M.S.L., S.R.R., B.T., L.L., I.W.B., E.C.M., K.S.C., and A.R. performed the experiments and analysis. P.D.G., D.A.F., W.W.M., D.J.W., and R.L.H. provided project supervision. All authors contributed to writing and editing the manuscript.

## Declaration of Interests

The authors declare no competing interests.

## STAR Methods

### Overview

All procedures were approved by the Institutional Animal Care and Use Committee of QTest Labs (Columbus, OH). Adult male Sprague Dawley rats (∼400-750 gm; N = 14) used in this study were housed one per cage (12 hr light/dark cycle; *ad libitum* access to food and water). The general aims of the study were to: 1) establish a model of myocardial ischemia, 2) utilize machine learning approaches to decode cardiovascular state changes, 3) determine if responsive closed-loop vagus nerve stimulation (VNS) controlled by an artificial neural network can significantly mitigate spontaneous myocardial ischemia, and 4) assess machine learning architectures for enhancing interpretability and facilitate detection of new emerging cardiovascular states. To acquire cardiovascular data, we recorded a lead II electrocardiogram (ECG), arterial blood pressure, and a photoplethysmogram (PPG) (schematic of experimental interfaces: Supplemental Fig. S1A). Analyses were performed in either MATLAB or Python.

### Surgery &Interface Placement

Animals were first administered Carprofen (5 mg/kg, s.c. injection) and anesthetized using isoflurane, similar to previous studies assessing VNS effects on cardiovascular physiology (Plachta et al., 2013 &2014). Isoflurane was vaporized into oxygen at 1.3-1.7%, and administered via a tracheotomy interface (Supplemental Fig. S1A, light red tube). Animals were kept supine throughout the procedure. Core body temperature was maintained at ∼37° C using a heating platform placed under the animal (Vestavia Scientific; Birmingham, AL).

The following 6 interfaces were next placed (schematic: Supplemental Fig. S1A): catheters were placed within the 1) right and 2) left femoral veins for intravenous (i.v.) administration of dobutamine and / or norepinephrine (see Inducing Myocardial Demand Ischemia Via Drug Injection for more details on drug administration); 3) arterial blood pressure (aBP) was recorded within the right carotid artery using a solid state blood pressure catheter (2 french; SPR-407 Mikro- Tip; Millar, Houston, Texas) and sent to a blood pressure amplifier (DA100C; BIOPAC, Goleta, CA); 4) a lead II electrocardiogram (ECG) was recorded using 3 hydrogel electrode contacts (ground: left arm, V+: right arm, V-: left leg) connected to an ECG amplifier (ECG100C; BIOPAC, Goleta, CA); 5) blood oxygen saturation level (SpO^2^) was recorded from the right hindpaw (OXY200; BIOPAC, Goleta, CA); 6) the left cervical vagus nerve was interfaced with a bipolar platinum iridium cuff electrode for delivering VNS, similar to our previous studies (Meyers et al., 2019; Ganzer et al., 2018). The bipolar VNS cuff electrode was tethered to a digitally controlled stimulator (Digitimer DS5; Hertfordshire, UK). Importantly, all instruments and stimulators were robustly electrically isolated to prevent stimulation artifact during cardiovascular data recordings.

### Vagus Nerve Stimulation (VNS) Cuff Implant

We interfaced with the left cervical vagus nerve to enable cardiovascular control, similar to several previous preclinical (Sachdeva et al., 2020; Plachta et al, 2013 &2014; Yamakawa et al., 2014; Shinlapawittayatorn et al., 2013) and human studies (Lewis et al., 2001; Anand et al., 2020). VNS was delivered with the following stimulation parameters: biphasic square wave morphology, 2-2.5 mA, 300 micro-second pulse width, at 30 Hz. VNS was delivered during closed-loop or open-loop stimulation regimes (see Modes of VNS Delivery for more details). Importantly, these VNS parameters are similar to previous preclinical studies using VNS for cardiovascular control (Sachdeva et al., 2020; Plachta et al, 2013 &2014; Yamakawa et al., 2014; Shinlapawittayatorn et al., 2013), and fall within clinically relevant stimulation ranges used in previous human trials using VNS for cardiovascular therapy (Anand et al., 2020; Table 1: Radcliffe et al., 2020).

### System Control for Signal Recording, Stimulation, and Injections

A schematic of the experiment and interfaces are shown in Supplemental Fig. S1A. All data was collected using a National Instruments USB-6259 data acquisition system (DAQ). The DAQ was controlled using MATLAB 2019a via a custom graphical user interface (The MathWorks; Natick, MA). We recorded 5 signals during the experiments: 1) the voltage sent to the VNS cuff electrodes, 2) the current drawn from the VNS cuff electrodes, 3) the lead II ECG waveform, 4) the aBP waveform, and 5) the SpO^2^ signal. Signals #1 and #2 were only active during VNS events. We also enabled 3 outputs during the experiments, as needed: 1 &2) triggers controlling the two drug injection pumps (KDS-200; Kent Scientific, Holliston, MA), and 3) a trigger controlling the VNS module. The DAQ operated at 10 kHz. This rate was needed to create VNS trains with the appropriate waveform morphology (e.g., biphasic square waves with the appropriate shape and resolution). Recorded signals were down-sampled and conditioned on-line as needed.

### Inducing Myocardial Ischemia Via Catecholamine Agent Injection

Cardiovascular stress and myocardial ischemia were induced using i.v. injection of dobutamine (∼2 μg x kg^-1^ x min^-1^) and / or norepinephrine (∼2 μg x kg^-1^ x min^-1^). Pilot studies were performed to assess dose dependent effects. These agents and similar dose rates have been used in several previous studies (Vimercati et al., 2012; Zhang &Mazgalev, 2009; Mandapaka &Hundley, 2006; Berk et a., 1977; Heusch &Ross, 1991).

### On-Line Cardiovascular Signal Conditioning and Feature Extraction

A schematic of the feature extraction is shown in Fig. 2. The subcomponents of features were first extracted online via the following signal conditioning processes (occurring every 100 ms): 1) a 10 kHz sampled epoch of the ECG and aBP waveforms were first down-sampled to 500 Hz sampled waveforms; 2) for the ECG epoch (Fig. 2A), the R waves were first detected using the ‘Peak Prominence’ attribute of the ‘findpeaks’ function in MATLAB 2019a. Window based detection was then used to identify the P, Ta, S, and T wave correlates (Fig. 2A). The time and voltage level of the given ECG wave correlates were recorded; 3) for the given aBP epoch (Fig. 2B), the systolic and diastolic pressure wave points were detected using the ‘Peak Prominence’ attribute of the ‘findpeaks’ function in MATLAB 2019a. The mmHg values of the systolic and diastolic aBP levels were recorded; 4) inhalation and exhalation cycles were encoded into the low frequency components of the aBP waveform. The linear envelope of the aBP waveform was calculated to extract the respiratory cycles time series. Inhalation points (i.e., breaths) were detected and recorded using the ‘Peak Prominence’ attribute of the ‘findpeaks’ function in MATLAB 2019a.

The thirteen-element feature vector was finally constructed from the above ECG and aBP waveform attributes via the following calculations (again, occurring every 100 ms):

- <u>Feature #1:</u> Heart Rate (beats per minute, or bpm) = R-R interval (s) / 60 s
- <u>Feature #2:</u> QRS Duration (ms) = relative Q wave to S wave duration
- <u>Feature #3:</u> RT Duration (ms) = R wave to T wave duration
- <u>Feature #4:</u> ST Duration (ms) = S wave to T wave duration
- <u>Feature #5:</u> Ta Level (mV) = voltage level of the Ta wave + voltage level of the TP interval
- <u>Feature #6:</u> R Level (mV) = voltage level of the R wave + voltage level of the TP interval
- <u>Feature #7:</u> ST Epoch Level (mV) = voltage level of the S wave + voltage level of the TP interval
- <u>Feature #8:</u> ST Slope (mV / s) = (T wave level (mV) – S wave level (mV)) / (T wave time (s) – S wave time (s))
- <u>Feature #9:</u> Diastolic Pressure (mmHg) = minimum pressure level during diastole
- <u>Feature #10:</u> Systolic Pressure (mmHg) = maximum pressure level during systole
- <u>Feature #11:</u> Mean Arterial Pressure (mmHg) = (systolic pressure (mmHg) + diastolic pressure (mmHg)) / 2
- <u>Feature #12:</u> Pulse Pressure (mmHg) = systolic pressure (mmHg) – diastolic pressure (mmHg)
- <u>Feature #13:</u> Breath Rate (breath rate per minute, or brpm) = breath count / 60 s

This feature vector contains relatively simple features that enhance decoding interpretation, and can be extracted for decoding without the need for burdensome compute power. To smooth the data, the feature vector was calculated every 100 ms and averaged over a 4 s sliding window continuously during real-time recordings. The feature data was recorded for offline analysis and was also used for online decoding *in vivo* (see Decoding Myocardial Demand Ischemia Using an Artificial Neural Network for more details).

### Decoding Myocardial Demand Ischemia Using an Artificial Neural Network (ANN)

#### Overview

A schematic of the artificial neural network (ANN) architecture and decoder outputs are shown in Supplemental Fig. S3A &S3B. We employed an ANN architecture and a supervised learning approach to decode 4 different cardiovascular states (i.e., classes): 1) rest (i.e., no drug injected), 2) dobutamine injection (D), 3) norepinephrine injection (NE), and 4) a combined dobutamine and norepinephrine injection (D+NE). The decoder outputs were assessed both offline and online to evaluate algorithm performance. Online predictions were used to either validate the ANN model or control closed-loop VNS.

#### Recording Events and Data Labels

Each recording contained the following events in sequence: 1) time 0 s = start of initial data streaming; 2) time 4 s = initiation of 4 s sliding window used for averaging features (sliding window increment per observation = 100 ms); 3) time 34 s = initiation of decoding (allows for 30 s of background feature data; this background feature data is used to both baseline subtract and standardize the subsequent recorded feature data); 4) time 94 s = start of a given injection; 5) time 214 s = end of injection, decoding, and recording.

To determine data labeling time points for supervised learning and ANN architecture attributes, we initially recorded pilot data from N=5 animals. On average, all 13 features statistically changed from baseline levels ∼15 s after an injection is started (feature changes were averaged across all 3 injection types). Said differently, average physiological changes across injection types occurred at 109 s. Therefore, the ‘rest’ class label occurred from 4 – 109 s, and the given drug’s class label occurred during the injection period from 109 - 214 s. This labeling approach enabled both physiological motivated data labels, and balanced durations of rest and a given cardiovascular stress state during a given recording (for mitigating class imbalance).

#### Grid Search for ANN Architecture and Hyperparameters

We used a grid search to arrive at an ANN architecture and hyperparameters (schematic of the final ANN architecture: Supplemental Fig. S3A). The grid search leveraged the same pilot data that was used for creating data labels described above (again from N=5 animals; a total of data ∼1.2 million points), and was performed on a computer with a graphical processing unit (NVIDIA GeForce GTX 1080; Santa Clara, CA). Overall accuracy (i.e., average accuracy across all classes) was used as the given algorithm’s performance metric. Our preliminary analysis demonstrated best performance using 2 hidden layers (a dense layer followed by a long short-term memory (LSTM) layer).

We next assessed combinations of the following architecture and hyperparameter values: 1) number of units in the dense layer (100, 250, or 500), 2) number of units in the LSTM layer (100, 250, or 500), 3) drop-out layer mask (between both the dense and LSTM and the LSTM and output layers; at 25%, 50%, or 75%), 4) mini-batch size (25%, 50%, or 75% of total data), and 6) early stopping criteria (reaching either 95% or 98% overall accuracy during training). The following were fixed during the grid search: sequence input layer size (13 units), output layer size (4 units), optimization algorithm (Adam), gradient decay metric (0.8), learning rate (0.01), gradient threshold (2), and L2 regularization metric (0.0005). The following ANN architecture and hyperparameters consistently performed the best, and were used throughout the study: <u>architecture</u> = sequence input layer (13 units), dense layer (250 units), drop-out layer mask (50%), LSTM layer (100 units), drop-out layer mask (25%), output layer (4 units); <u>hyperparameters</u> = mini-batch size (75%), early stopping (reaching 98% overall accuracy during training).

## Algorithm Performance Evaluation

### Online In Vivo Assessments (related to Fig. 5 and Supplemental Fig. S5)

We performed online *in vivo* decoding of cardiovascular state and ANN controlled VNS in a total of N=9 animals. Overall, we modeled a clinical use case for the ANN. We continuously added to the base training set across experiments, and performed supervised updating of the ANN within a given animal for subsequent real-time prediction and closed-loop ANN-VNS control. <u>Experimental design details</u>: 1) the initial training set consisted of the pilot data; 2) each subsequent new animal then contributed 6 more recordings to the base training set (injection order randomized; 2 injections of D, 2 injections of NE, and 2 injections of D+NE); 3) for a given new animal, a new ANN model was trained and validated online *in vivo*; 4) online testing consisted of real time prediction during 3 injections of the target ischemic state (D+NE). We report overall accuracy for online *in vivo* ANN performance (Supplemental Fig. S5A).

### Offline Assessments (*related to Fig. 4, Supplemental Fig. S3, and Supplemental Fig. S4*)

We also assessed algorithm performance offline using the final data set (pilot data [N=5] + experimental data [N=9]). 10-fold cross validation was used to appraise the performance of the ANN and other types of classifiers for comparison (using 80% / 20% train / test splits, respectively). We compared the ANN performance to 3 other classifier types: 1) an ANN-NO-LSTM architecture (via replacing the LSTM layer with a second hidden dense layer); 2) a support vector machine (SVM); and 3) a linear discriminant analysis (LDA). Similar to the ANN, we performed a hyperparameter grid search to optimize the performance of both the SVM and LDA. We report overall accuracy for all classification approaches (Supplemental Fig. S3C).

### Modes of VNS Delivery (related to Fig. 5, Supplemental Fig. S5, and Supplemental Fig. S6)

We assessed the effects of ANN-VNS during episodes of spontaneous myocardial ischemia (i.e., ‘Isch.’ induced by D+NE). Supplemental Fig. S5B shows the average ANN decoder outputs during real-time predictions *in vivo* (decoder outputs averaged across all N=9 animals). Closed-loop VNS was triggered when the ‘Isch.’ decoder output score was greater than 0.5, representing a class probability greater than 50% (similar to our previous decoding and device activation studies: Ganzer et al., 2020; Bouton et al., 2016). Once triggered by the ANN, VNS remained active for the remainder of the injection (i.e., up until 214 s). Importantly, all instrumentation and stimulators were robustly isolated to prevent stimulation artifact during recordings.

#### Additional VNS Controls

We performed 2 VNS controls, complimenting ANN-VNS. The first VNS control condition was open-loop VNS (data presented in Fig. 5 and Supplemental Fig. S6B). We used a 20% VNS ON / 80% VNS OFF duty cycle for the open-loop VNS condition, to model preprogrammed open-loop VNS duty cycles used in clinical trials for cardiovascular treatments (Anand et al., 2020; Table 1: Radcliffe et al., 2020). Open-loop VNS recordings lasted a total of ∼500 s, with 2-3 recording replicates within an animal (across N=5 animals). A D+NE injection was started at a randomized time during an open-loop VNS recording epoch, using the same 2 min injection duration.

The second VNS control condition was ANN-VNS following a vagotomy caudal to the cervical VNS site (data presented in Supplemental Fig. S6C). The vagus nerve was cut using surgical scissors and the cut ends were further separated by ∼1 mm to ensure a complete vagotomy. Recording and VNS was resumed approximately 30 mins after the vagotomy to allow for physiological equilibration. We performed 3 recording replicates within an animal (across N=3 animals).

### Data Analyses for Assessing Cardiovascular Feature Changes (With or Without VNS)

Several cardiovascular features shown throughout the manuscript are presented as a change from baseline (i.e., Δ relative to baseline). For the given feature, baseline activity from the first 30 s of a recording was used to create the baseline subtracted feature time series (see On-Line Cardiovascular Signal Conditioning and Feature Extraction for details on feature creation). A given feature was further processed to assess effects as follows:

- **Related to Fig. 1D:** Rate-pressure product (RPP) across time was not a component of the overall 13-element feature vector, but was calculated similar to previous studies (Gobel et al., 1978): RPP = (Heart Rate x Systolic Blood pressure) / 100.
- **Related to Fig. 1G:** Each point is a single animal’s recording for a given drug condition. For a given point, the Δ ST epoch level or Δ RPP value was its average during the entire injection period, relative to baseline. We report the Pearson’s correlation coefficient R.
- **Related to Fig. 3:** A given feature’s Δ value was its average during the entire injection period, relative to baseline.
- **Related to Supplemental Fig. S2A &S2B:** We compared the variability (i.e., entropy) of cardiovascular changes for our preclinical rat data and human data collected in other studies. We used the following human cardiovascular data acquired from the physionet.org database (Goldberger et al., 2000): Supplemental Fig. S2A, ‘Human 1’ = recorded in the intensive care unit (Kim et al., 2016); Supplemental Fig. S2B, ‘Human 2’ (Taddei et al., 1992) &‘Human 3’ (Jager et al., 2003) = recorded during ambulatory episodes of myocardial ischemia. The cardiovascular feature matrices were next prepared using either all 13 features (Supplemental Fig. S2A) or only the 8 ECG features (Supplemental Fig. S2B). The feature matrices for the human data used a modified version of the rat feature extraction algorithm. Entropy was finally calculated and reported using the ‘entropy’ function in MATLAB 2019a.
- **Related to Supplemental Fig. S2C:** We assessed the relationship between different pairs of drug states and report the Pearson’s correlation coefficient R. Within each animal (N=9), we calculate the R values for all possible pairs of injections using the feature matrix across time (e.g., a D+NE &NE correlation). We plot each animal’s average R (single points in the figure) across the 3 different types of injection correlations.
- **Related to Fig. 5C, 5D, and Supplemental Fig. S6:** In Fig. 5D, we report two additional features: SpO^2^ = blood oxygen saturation value from the PPG monitor, and the QT / TQ ratio (related to arrythmia probability; Fossa et al., 2017; relevant wave point correlates are shown in Fig. 2). Overall, for ‘Isch.’ a given feature’s Δ value was its average during the entire injection period, relative to baseline. Overall, for ‘Isch. + ANN-VNS’, ‘Isch. + OL-VNS’, and ‘Isch. + ANN-VNS + (xcv)’ a given feature’s Δ value was its average while VNS was on, relative to baseline.

### Detecting New Emerging Cardiovascular Stress States (related to Fig. 6 and Supplemental Fig. S7)

In a subset of animals (N=4), we recorded the 13 features during injections at a higher dose rate (10 μg x kg^-1^ x min^-1^) across 3 injection types: ‘high dose dobutamine’ = H-D; ‘high dose norepinephrine’ = H-NE; ‘high dose dobutamine &norepinephrine combined’ = H-D+NE. These recordings at a ∼5x higher dose rate presented a significantly different feature profile during a given injection (data not shown) and were used for subsequent emerging stress state detection. We appraised the ability of 3 techniques to detect these emerging stress states (emerging state / outlier detection technique review: Park, 2019): 1) LSTM autoencoder (LSTM-AE), 2) sparse autoencoder (Sparse-AE), and 3) isolation forest (Iso-Forest). Each technique was optimized using a grid search on a subset of the data (LSTM-AE major parameters: encoder layer (input) = 2730 units, hidden layer = 256 units, decoder layer (output) = 2730 units, L2 regularization = 0; Sparse-AE major parameters: encoder layer (input) = 546 units, hidden layer = 50 units, decoder layer (output) = 546 units, L2 regularization = 0.1, sparsity proportion = 1; Iso-Forest major parameters: N estimators = 2, max features = 140, outlier proportion = 0.16). We finally performed a 10-fold cross validation to appraise the performance of the 3 techniques (using 80% / 20% train / test splits, respectively). We report ‘emerging stress state detection sensitivity’ for the 3 techniques (Fig. 6; i.e., true positive rate when presented with a new emerging stress state, averaged across the 3 types of emerging stress states).

### Visualizing the ‘Cardiovascular Latent Space’ Using LSTM Autoencoders &UMAP (related to Fig. 7A, 7B, Supplemental Video 1, and Supplemental Fig. S8)

We used the hidden layer of the LSTM-AE combined with the dimensionality reduction method uniform manifold approximation and projection (or UMAP; McInnes &Healy, 2018) to visualize interpretable 2-dimensional representations of the emerging stress states (i.e., the ‘cardiovascular latent space’). The hidden layer of autoencoders and UMAP are both commonly used for generating latent features and dimensionality reduction (McConville et al., 2019). The hidden layer of the LSTM-AE (256 elements) was labeled and passed into the UMAP algorithm for supervised dimensionality reduction (schematic of architecture: Fig. 7A; final UMAP hyperparameters: n_neighbors = 15; min_dist = 0.1; n_components = 2). The ability to separate the 2-dimensional known and emerging stress state points in the ‘cardiovascular latent space’ was assessed using a linear SVM (related to Supplemental Fig. S8).

### Identifying New Emerging Cardiovascular Stress State Types Using Unsupervised Clustering (related to Fig. 7C)

We performed unsupervised clustering of the 2-dimensional ‘cardiovascular latent space’ points using the hierarchical density based spatial clustering of applications with noise (HDBSCAN) method. HDBSCAN is a robust unsupervised clustering technique that deals well with diverse clustering scenarios. For a given stress state presentation scenario (combinations of 1, 2, or 3 emerging stress types), the unsupervised HDBSCAN method was first challenged to cluster the 2-dimensional ‘cardiovascular latent space’ points (i.e., determine the number of emerging stress states present). We next calculated clustering performance (V-measure * 100%, Rosenberg &Hirschberg, 2007; V-measure is a well-studied metric for assessing clustering quality; 0 = completely random clustering, 1 = perfect clustering). In this implementation, V-measure based performance also quantifies correlates of several processes including the LSTM- AE’s ability to generate useful latent space vectors, the quality of the subsequent UMAP dimensionality reduction, and the quality of the final HDBSCAN method. We calculate and report performance (Fig. 7C) averaged across the 7 stress state presentation scenarios (H-D alone, H-NE alone, H-D+NE alone, H-D &H-NE, H-D &H-D+NE, H-NE &H-D+NE, and H-D &H-NE &H-D+NE).

### Statistics

Normality tests were performed for each analysis to determine if parametric or nonparametric statistics should be used. All statistical tests were two-tailed unless otherwise noted, and were performed in GraphPad Prism. An alpha level of 0.05 was accepted for significance, unless Bonferroni corrections are noted. Chance performance levels were generated by randomly permuting the true data labels 10 times, similar to previous studies (Ganzer et al., 2020; Ojala &Garriga, 2010).

We report the Pearson’s correlation coefficient R for data in panel Fig. 1G and Supplemental Fig. S2C. Effects of injections on the 13-element feature vector were evaluated using a separate one-way ANOVA for each feature (related Fig. 3). The factor was injection type with 3 levels: D, NE, and D+NE. Tukey’s post-hoc test was used to determine differences for a given feature across injection types.

Differences in classification performance were evaluated using a one-way ANOVA (related Supplemental Fig. S3C). The factor was classifier type with 4 levels: ANN, ANN-NO-LSTM, SVM, and LDA. Tukey’s post-hoc test was used to determine differences in performance across classifier types. A one-tailed independent samples t-test was used to determine if ANN performance values were above chance levels (related to Fig. 4D, confusion matrix). A Bonferroni corrected alpha value of 0.003 was used for significance (0.05 / 16 comparisons).

Differences in cardiovascular biomarkers were assessed using separate one-way ANOVAs for each main effect (related to Fig. 5C) and side effect (related to Fig. 5D). The factor was state type with 3 levels: Isch., Isch. + ANN-VNS, and Isch. + OL-VNS. Tukey’s post-hoc test was used to determine differences across state types. VNS ON time was also assessed using a one-way ANOVA (related to Supplemental Fig. S5).

2-dimensional known and emerging stress state points in the ‘cardiovascular latent space’ were separated using a linear boundary (related to Supplemental Fig. S8), and differences between actual and chance performance were assessed using a t-test. Differences in emerging stress state detection performance were evaluated using a one-way ANOVA (related Fig. 6A). The factor was detection technique with 3 levels: LSTM-AE, Sparse-AE, and Iso-Forest. Tukey’s post-hoc test was used to determine differences in performance across detection techniques. A receiver operating characteristic curve was generated for the LSTM-AE approach to present the raw MSE data (related to Supplemental Fig. S7). Lastly, we report the performance for unsupervised clustering and detection of different types of emerging stress states (related to Fig. 7C; performance = V-measure * 100%; Rosenberg &Hirschberg, 2007). We assessed differences between actual and chance performance using a t-test (either averaged across the emerging stress state presentation scenarios = Fig. 7C; or assessed separately across the emerging stress state presentation scenarios = Supplemental Fig. S9).

## Notes

### Competing Interest Statement

The authors have declared no competing interest.

## References

Global Health Estimates 2016: Deaths by Cause, Age, Sex, by Country and by Region, 2000-2016. Geneva, World Health Organization; 2018.

Ardehali, A., & Ports, T. A. (1990). Myocardial oxygen supply and demand. Chest, 98(3), 699–705.

Deedwania, P. C., & Carbajal, E. V. (1992). Role of myocardial oxygen demand in the pathogenesis of silent ischemia during daily life. The American journal of cardiology, 70(16), F19–F24.

Hinderliter, A., Miller, P., Bragdon, E., Ballenger, M., & Sheps, D. (1991). Myocardial ischemia during daily activities: the importance of increased myocardial oxygen demand. Journal of the American College of Cardiology, 18(2), 405–412.

Braun, M. M., Stevens, W. A., & Barstow, C. (2018). Stable coronary artery disease: treatment. American family physician, 97(6), 376–384.

Conti, C. R., Bavry, A. A., & Petersen, J. W. (2012). Silent ischemia: clinical relevance. Journal of the American College of Cardiology, 59(5), 435–441.

Cohn, P. F. (1998). Treatment of chronic myocardial ischemia: rationale and treatment options. Cardiovascular drugs and therapy, 12(3), 217–223.

Gutterman, D. D. (2009). Silent myocardial ischemia. Circulation Journal, 73(5), 785–797.

Cecchi, A. C., Dovellini, E. V., Marchi, F., Pucci, P., Santoro, G. M., & Fazzini, P. F. (1983). Silent myocardial ischemia during ambulatory electrocardiographic monitoring in patients with effort angina. Journal of the American College of Cardiology, 1(3), 934–939.

Rozanski A, Berman DS: Silent myocardial ischemia.I. Pathophysiology, frequency of occurrence, and approaches toward detection. Am Heart J 114:615, 1987.

Schwartz, B. G., Kloner, R. A., & Naghavi, M. (2018). Acute and subacute triggers of cardiovascular events. The American journal of cardiology, 122(12), 2157–2165.

Capilupi, M. J., Kerath, S. M., & Becker, L. B. (2020). Vagus nerve stimulation and the cardiovascular system. Cold Spring Harbor Perspectives in Medicine, 10(2), a034173.

Levy MN, Schwartz PJ, eds. Vagal Control of the Heart: Experimental Basis and Clinical Implications. Armonk, NY: Futura Publishing Co; 1994:644

Ardell, J. L., Rajendran, P. S., Nier, H. A., Kenknight, B. H., & Armour, J. A. (2015). Central-peripheral neural network interactions evoked by vagus nerve stimulation: functional consequences on control of cardiac function. American Journal of Physiology-Heart and Circulatory Physiology, 309(10), H1740–H1752.

Buck, J. D., Warltier, D. C., Hardman, H. F., & Gross, G. J. (1981). Effects of sotalol and vagal stimulation on ischemic myocardial blood flow distribution in the canine heart. Journal of Pharmacology and Experimental Therapeutics, 216(2), 347–351.

Patel, D. J., Mulcahy, D., Norrie, J., Wright, C., Clarke, D., Ford, I., & Fox, K. M. (1996). Natural variability of transient myocardial ischaemia during daily life: an obstacle when assessing efficacy of anti-ischaemic agents?. Heart, 76(6), 477–482.

Celermajer, D. S., Spiegelhalter, D. J., Deanfield, M., & Deanfield, J. E. (1994). Variability of episodic ST segment depression in chronic stable angina: implications for individual and group trials of therapeutic efficacy. Journal of the American College of Cardiology, 23(1), 66–73.

Deanfield, J. E., & Spiegelhalter, D. (1990). Variability of myocardial ischemia: implications for monitoring strategies. In Silent Myocardial Ischemia: A Critical Appraisal (Vol. 37, pp. 176–186). Karger Publishers.

Tzivoni, D., Gavish, A., Benhorin, J., Banai, S., Keren, A., & Stern, S. (1987). Day-to-day variability of myocardial ischemic episodes in coronary artery disease. The American journal of cardiology, 60(13), 1003–1005.

Michaelides, A. P., Liakos, C. I., Antoniades, C., Tsiachris, D. L., Soulis, D., Dilaveris, P. E., … & Stefanadis, C. I. (2010). ST-segment depression in hyperventilation indicates a false positive exercise test in patients with mitral valve prolapse. Cardiology research and practice, 2010.

Petrov, D. B., Sardovski, S. I., & Milanova, M. H. (2012). Severe hypokalemia masquerading myocardial ischemia. Cardiology Research, 3(5), 236.

Sapin, P. M., Koch, G., Blauwet, M. B., Mccarthy, J. J., Hinds, S. W., & Gettes, L. S. (1991). Identification of false positive exercise tests with use of electrocardiographic criteria: a possible role for atrial repolarization waves. Journal of the American College of Cardiology, 18(1), 127–135.

Wright, J., Macefield, V. G., Van Schaik, A., & Tapson, J. C. (2016). A review of control strategies in closed-loop neuroprosthetic systems. Frontiers in neuroscience, 10, 312.

Sun, F. T., & Morrell, M. J. (2014). Closed-loop neurostimulation: the clinical experience. Neurotherapeutics, 11(3), 553–563.

Hays SA (2016) Enhancing rehabilitative therapies with vagus nerve stimulation. Neurotherapeutics 13:382–394.

Ganzer PD, Darrow MJ, Meyers EC, Solorzano BR, Ruiz AD, Robertson NM, Adcock KS, James JT, Jeong HS, Becker AM, Goldberg MP, Pruitt DT, Hays SA, Kilgard MP, Rennaker RL 2nd (2018) Closed-loop neuromodulation restores network connectivity and motor control after spinal cord injury. Elife 7:e32058.

Ganzer, P. D., & Sharma, G. (2019). Opportunities and challenges for developing closed-loop bioelectronic medicines. Neural Regeneration Research, 14(1), 46.

Vellido, A. (2019). The importance of interpretability and visualization in machine learning for applications in medicine and health care. Neural Computing and Applications, 1–15.

Tonekaboni, S., Joshi, S., Mccradden, M. D., & Goldenberg, A. (2019). What clinicians want: contextualizing explainable machine learning for clinical end use. arXiv preprint arXiv:1905.05134.

Tjoa, E., & Guan, C. (2019). A survey on explainable artificial intelligence (XAI): towards medical XAI. arXiv preprint arXiv:1907.07374.

Liu S, Wang X, Liu M, Zhu J (2017) Towards better analysis of machine learning models: a visual analytics perspective. Vis Inf 1(1):48–56.

Vellido, A., MartíN-Guerrero, J. D., & Lisboa, P. J. (2012). Making machine learning models interpretable. In ESANN (Vol. 12, pp. 163–172).

Vellido Alcacena, A., MartíN, J. D., Rossi, F., & Lisboa, P. J. (2011). Seeing is believing: The importance of visualization in real-world machine learning applications. In Proceedings: 19th European Symposium on Artificial Neural Networks, Computational Intelligence and Machine Learning, ESANN 2011: Bruges, Belgium, April 27-28-29, 2011 (pp. 219–226).

Zahavy, T., Ben-Zrihem, N., & Mannor, S. (2016, June). Graying the black box: Understanding dqns. In International Conference on Machine Learning (pp. 1899–1908).

Epel, E. S., Crosswell, A. D., Mayer, S. E., Prather, A. A., Slavich, G. M., Puterman, E., & Mendes, W. B. (2018). More than a feeling: A unified view of stress measurement for population science. Frontiers in neuroendocrinology, 49, 146–169.

Rocco MB, Nabel EG, Selwyn AP. Development and validation of ambulatory monitoring to characterize ischemic heart disease out of hospital. Cardiology Clinics 1986; 4: 659 – 668.

Rehman, A., Zalos, G., Andrews, N. P., Mulcahy, D., & Quyyumi, A. A. (1997). Blood pressure changes during transient myocardial ischemia: insights into mechanisms. Journal of the American College of Cardiology, 30(5), 1249–1255.

Deedwania, P. C., & Nelson, J. R. (1990). Pathophysiology of silent myocardial ischemia during daily life. Hemodynamic evaluation by simultaneous electrocardiographic and blood pressure monitoring. Circulation, 82(4), 1296–1304.

Mandapaka, S., & Hundley, W. G. (2006). Dobutamine cardiovascular magnetic resonance: a review. Journal of Magnetic Resonance Imaging: An Official Journal of the International Society for Magnetic Resonance in Medicine, 24(3), 499–512.

Heusch G & Ross J (ed.) (1991) Adrenergic mechanisms in myocardial ischemia. Darmstadt: Steinkopff; New York: Springer.

Kawada, T., Yamazaki, T., Akiyama, T., Mori, H., Inagaki, M., Shishido, T., … & Sunagawa, K. (2002). Effects of brief ischaemia on myocardial acetylcholine and noradrenaline levels in anaesthetized cats. Autonomic Neuroscience, 95(1-2), 37–42.

Barger, A. C., Herd, J. A., & Liebowitz, M. R. (1961). Chronic Catheterization of Coronary Artery: Induction of ECG Pattern of Myocardial Ischemia by Intracoronary Epinephrine. Proceedings of the Society for Experimental Biology and Medicine, 107(3), 474–477.

Lepeschkin, E., Marchet, H., Schroeder, G., Wagner, R., E Silva, P. D. P., & Raab, W. (1960). Effect of epinephrine and norepinephrine on the electrocardiogram of 100 normal subjects∗. The American Journal of Cardiology, 5(5), 594–603.

Segar, D. S., Ryan, T., Sawada, S. G., Johnson, M., & Feigenbaum, H. (1995). Pharmacologically induced myocardial ischemia: a comparison of dobutamine and dipyridamole. Journal of the American Society of Echocardiography, 8(1), 9–14.

Detry, J. M. R., Piette, F., & Brasseur, L. A. (1970). Hemodynamic determinants of exercise ST-segment depression in coronary patients. Circulation, 42(4), 593–599.

Gobel, F. L., Norstrom, L. A., Nelson, R. R., Jorgensen, C. R., & Wang, Y. (1978). The rate-pressure product as an index of myocardial oxygen consumption during exercise in patients with angina pectoris. Circulation, 57(3), 549–556.

Klabunde, R. E. (2017). Cardiac electrophysiology: normal and ischemic ionic currents and the ECG. Advances in physiology education, 41(1), 29–37.

Janse, M. J. (2007). ST-segment elevation or TQ-segment depression?. Heart rhythm, 4(2), 207.

KléBer, A. G., Janse, M. J., Van Capelle, F. J., & Durrer, D. I. R. K. (1978). Mechanism and time course of ST and TQ segment changes during acute regional myocardial ischemia in the pig heart determined by extracellular and intracellular recordings. Circulation Research, 42(5), 603–613.

Cinca, J., Janse, M. J., MoréNa, H., Candell, J., Valle, V., & Durrer, D. (1980). Mechanism and time course of the early electrical changes during acute coronary artery occlusion: an attempt to correlate the early ECG changes in man to the cellular electrophysiology in the pig. Chest, 77(4), 499–505.

Sharma, N., & Gedeon, T. (2012). Objective measures, sensors and computational techniques for stress recognition and classification: A survey. Computer methods and programs in biomedicine, 108(3), 1287–1301.

Goldberger, A. L., Amaral, L. A., Glass, L., Hausdorff, J. M., Ivanov, P. C., Mark, R. G., … & Stanley, H. E. (2000). PhysioBank, PhysioToolkit, and PhysioNet: components of a new research resource for complex physiologic signals. circulation, 101(23), e215–e220.

Kim, N., Krasner, A., Kosinski, C., Wininger, M., Qadri, M., Kappus, Z., … & Craelius, W. (2016). Trending autoregulatory indices during treatment for traumatic brain injury. Journal of clinical monitoring and computing, 30(6), 821–831.

Taddei, A., Distante, G., Emdin, M., Pisani, P., Moody, G. B., Zeelenberg, C., & Marchesi, C. (1992). The European ST-T database: standard for evaluating systems for the analysis of ST-T changes in ambulatory electrocardiography. European heart journal, 13(9), 1164–1172.

Jager, F., Taddei, A., Moody, G. B., Emdin, M., Antolič, G., Dorn, R., … & Mark, R. G. (2003). Long-term ST database: a reference for the development and evaluation of automated ischaemia detectors and for the study of the dynamics of myocardial ischaemia. Medical and Biological Engineering and Computing, 41(2), 172–182.

Murat, F., Yildirim, O., Talo, M., Baloglu, U. B., Demir, Y., & Acharya, U. R. (2020). Application of deep learning techniques for heartbeats detection using ECG signals-analysis and review. Computers in Biology and Medicine, 103726.

Gers, F. A., Schmidhuber, J., & Cummins, F. (1999). Learning to forget: Continual prediction with LSTM.

Svensson, P., Niklasson, U., & ÖStergren, J. (2001). Episodes of ST‐segment depression is related to changes in ambulatory blood pressure and heart rate in intermittent claudication. Journal of internal medicine, 250(5), 398–405.

Fossa, A. A. (2017). Beat‐to‐beat ECG restitution: A review and proposal for a new biomarker to assess cardiac stress and ventricular tachyarrhythmia vulnerability. Annals of Noninvasive Electrocardiology, 22(5), e12460.

Anand, I. S., Konstam, M. A., Klein, H. U., Mann, D. L., Ardell, J. L., Gregory, D. D., … & Butler, J. (2020). Comparison of symptomatic and functional responses to vagus nerve stimulation in ANTHEM‐HF. INOVATE‐HF, and NECTAR‐HF. ESC heart failure, 7(1), 76–84.

Radcliffe, E. J., Pearman, C. M., Watkins, A., Lawless, M., Kirkwood, G. J., Saxton, S. N., … & Trafford, A. W. (2020). Chronic vagal nerve stimulation has no effect on tachycardia‐induced heart failure progression or excitation–contraction coupling. Physiological Reports, 8(2).

Park, C. H. (2019). Outlier and anomaly pattern detection on data streams. The Journal of Supercomputing, 75(9), 6118–6128.

Mcinnes, L., Healy, J., & Melville, J. (2018). Umap: Uniform manifold approximation and projection for dimension reduction. arXiv preprint arXiv:1802.03426.

Mcconville, R., Santos-Rodriguez, R., Piechocki, R. J., & Craddock, I. (2019). N2d:(not too) deep clustering via clustering the local manifold of an autoencoded embedding. arXiv preprint arXiv:1908.05968.

Campello R.J.G.B., Moulavi D., Sander J. (2013) Density-Based Clustering Based on Hierarchical Density Estimates. In: Pei J., Tseng V.S., Cao L., Motoda H., Xu G. (eds) Advances in Knowledge Discovery and Data Mining. PAKDD 2013. Lecture Notes in Computer Science, vol 7819. Springer, Berlin, Heidelberg. https://doi.org/10.1007/978-3-642-37456-2_14

Rosenberg, A., & Hirschberg, J. (2007). V-measure: A conditional entropy-based external cluster evaluation measure. In Proceedings of the 2007 joint conference on empirical methods in natural language processing and computational natural language learning (EMNLP-CoNLL) (pp. 410–420).

Malliani A (1986). The elusive link between transient myocardial ischemia and pain. Circulation, 73(2), 201–204.

Soliman, E. Z. (2019). Silent myocardial infarction and risk of heart failure: current evidence and gaps in knowledge. Trends in cardiovascular medicine, 29(4), 239–244.

Ding, K., & Kullo, I. J. (2009). Evolutionary genetics of coronary heart disease. Circulation, 119(3), 459–467.

Kember, G., Armour, J. A., & Zamir, M. (2013). Neural control hierarchy of the heart has not evolved to deal with myocardial ischemia. Physiological genomics, 45(15), 638–644.

Swap, C. J., & Nagurney, J. T. (2005). Value and limitations of chest pain history in the evaluation of patients with suspected acute coronary syndromes. Jama, 294(20), 2623–2629.

Pomblum, V. J., Korbmacher, B., Cleveland, S., Sunderdiek, U., Klocke, R. C., & Schipke, J. D. (2010). Cardiac stunning in the clinic: the full picture. Interactive cardiovascular and thoracic surgery, 10(1), 86–91.

Tabibiazar, R., & Edelman, S. V. (2003). Silent ischemia in people with diabetes: a condition that must be heard. Clinical Diabetes, 21(1), 5–9.

Pop-Busui, R. (2010). Cardiac autonomic neuropathy in diabetes: a clinical perspective. Diabetes care, 33(2), 434–441.

Hassabis, D., Kumaran, D., Summerfield, C., & Botvinick, M. (2017). Neuroscience-inspired artificial intelligence. Neuron, 95(2), 245–258.

Marblestone, A. H., Wayne, G., & Kording, K. P. (2016). Toward an integration of deep learning and neuroscience. Frontiers in computational neuroscience, 10, 94.

Wilson, H. J., & Daugherty, P. R. (2018). Collaborative intelligence: humans and AI are joining forces. Harvard Business Review, 96(4), 114–123.

Jarrahi, M. H. (2018). Artificial intelligence and the future of work: Human-AI symbiosis in organizational decision making. Business Horizons, 61(4), 577–586.

Lotze, U., ÖZbek, C., Gerk, U., Kaufmann, H., Sen, S., & Figulla, H. R. (1999). Three-year follow-up of patients with silent ischemia in the subacute phase of myocardial infarction after thrombolysis and early coronary intervention. International journal of cardiology, 71(2), 167–178.

Deedwania, P. C., & Carbajal, E. V. (1990). Silent ischemia during daily life is an independent predictor of mortality in stable angina. Circulation, 81(3), 748–756.

Balla, C., Pavasini, R., & Ferrari, R. (2018). Treatment of angina: where are we?. Cardiology, 140(1), 52–67.

Vitale, F., & Litt, B. (2018). Bioelectronics: the promise of leveraging the body’s circuitry to treat disease.

Birmingham K, Gradinaru V, Anikeeva P, Grill WM, Pikov V, Mclaughlin B, Pasricha P, Weber D, Ludwig K, Famm K (2014) Bioelectronic medicines: a research roadmap. Nat Rev Drug Discov 13:399–400.

Machhada, A., Hosford, P. S., Dyson, A., Ackland, G. L., Mastitskaya, S., & Gourine, A. V. (2020). Optogenetic Stimulation of Vagal Efferent Activity Preserves Left Ventricular Function in Experimental Heart Failure. JACC: Basic to Translational Science.

Nuntaphum, W., Pongkan, W., Wongjaikam, S., Thummasorn, S., Tanajak, P., Khamseekaew, J., … & Shinlapawittayatorn, K. (2018). Vagus nerve stimulation exerts cardioprotection against myocardial ischemia/reperfusion injury predominantly through its efferent vagal fibers. Basic research in cardiology, 113(4), 22.

Del Rio, C. L., Dawson, T. A., Clymer, B. D., Paterson, D. J., & Billman, G. E. (2008). Effects of acute vagal nerve stimulation on the early passive electrical changes induced by myocardial ischaemia in dogs: heart rate‐mediated attenuation. Experimental physiology, 93(8), 931–944.

Vanoli, E., De Ferrari, G. M., Stramba-Badiale, M., Hull Jr, S. S., Foreman, R. D., & Schwartz, P. J. (1991). Vagal stimulation and prevention of sudden death in conscious dogs with a healed myocardial infarction. Circulation research, 68(5), 1471–1481.

Myers R. W., Pearlman A. S., Hyman R. M., Goldstein R. A., Kent K. M., Goldstein R. E., & Epstein S. E. (1974). Beneficial effects of vagal stimulation and bradycardia during experimental acute myocardial ischemia. Circulation, 49(5), 943–947.

Pepine, C. J., Geller, N. L., Knatterud, G. L., Bourassa, M. G., Chaitman, B. R., Davies, R. F., … & Mueller, H. (1994). The Asymptomatic Cardiac Ischemia Pilot (ACIP) study: design of a randomized clinical trial, baseline data and implications for a long-term outcome trial. Journal of the American College of Cardiology, 24(1), 1–10.

Trimarco, B., Chierchia, S., Lembo, G., De Luca, N., Ricciardelli, B., Condorelli, G., … & Condorelli, M. (1990). Prolonged duration of myocardial ischemia in patients with coronary heart disease and impaired cardiopulmonary baroreceptor sensitivity. Circulation, 81(6), 1792–1802.

Huang, W., Lai, K. K., Nakamori, Y., Wang, S., and Yu, L., 2007, “Neural Networks in Finance and Economics Forecasting,” Int. J. Info. Tech. Dec. Mak., 06(01), pp. 113–140.

Wang, M., Zhao, L., Du, R., Wang, C., Chen, L., Tian, L., and Eugene Stanley, H., 2018, “A Novel Hybrid Method of Forecasting Crude Oil Prices Using Complex Network Science and Artificial Intelligence Algorithms,” Applied Energy, 220, pp. 480–495.

Miller, D. D., and Brown, E. W., 2018, “Artificial Intelligence in Medical Practice: The Question to the Answer?,” The American Journal of Medicine, 131(2), pp. 129–133.

Choy, G., Khalilzadeh, O., Michalski, M., Do, S., Samir, A. E., Pianykh, O. S., Geis, J. R., Pandharipande, P. V., Brink, J. A., and Dreyer, K. J., 2018, “Current Applications and Future Impact of Machine Learning in Radiology,” Radiology, 288(2), pp. 318–328.

Romiti, S., Vinciguerra, M., Saade, W., Anso Cortajarena, I., & Greco, E. (2020). Artificial Intelligence (AI) and Cardiovascular Diseases: An Unexpected Alliance. Cardiology Research and Practice, 2020.

Kawada, T., & Sugimachi, M. (2009). Artificial neural interfaces for bionic cardiovascular treatments. Journal of Artificial Organs, 12(1), 17–22.

Gotoh, T. M., Tanaka, K., & Morita, H. (2005). Controlling arterial blood pressure using a computer–brain interface. Neuroreport, 16(4), 343–347.

Sugimachi, M., & Sunagawa, K. (2009). Bionic cardiology: Exploration into a wealth of controllable body parts in the cardiovascular system. IEEE reviews in biomedical engineering, 2, 172–186.

Sato, T., Kawada, T., Sugimachi, M., & Sunagawa, K. (2002). Bionic technology revitalizes native baroreflex function in rats with baroreflex failure. Circulation, 106(6), 730–734.

Jungmann, F., Jorg, T., Hahn, F., Pinto dos Santos, D., Jungmann, S. M., DüBer, C., Mildenberger, P., and Kloeckner, R., 2020, “Attitudes Toward Artificial Intelligence Among Radiologists, IT Specialists, and Industry,” Academic Radiology.

Holzinger, A., Langs, G., Denk, H., Zatloukal, K., and MüLler, H., 2019, “Causability and Explainability of Artificial Intelligence in Medicine,” WIREs Data Mining and Knowledge Discovery, 9(4), p. e1312.

Plachta, D. T., Gierthmuehlen, M., Cota, O., Boeser, F., & Stieglitz, T. (2013). BaroLoop: Using a multichannel cuff electrode and selective stimulation to reduce blood pressure. In 2013 35th Annual International Conference of the IEEE Engineering in Medicine and Biology Society (EMBC) (pp. 755-758). IEEE.

Plachta, D. T., Gierthmuehlen, M., Cota, O., Espinosa, N., Boeser, F., Herrera, T. C., … & Zentner, J. (2014). Blood pressure control with selective vagal nerve stimulation and minimal side effects. Journal of neural engineering, 11(3), 036011.

Ganzer, P. D., Darrow, M. J., Meyers, E. C., Solorzano, B. R., Ruiz, A. D., Robertson, N. M., … & Goldberg, M. P. (2018). Closed-loop neuromodulation restores network connectivity and motor control after spinal cord injury. Elife, 7, e32058.

Meyers, E. C., Kasliwal, N., Solorzano, B. R., Lai, E., Bendale, G., Berry, A., … & Hays, S. A. (2019). Enhancing plasticity in central networks improves motor and sensory recovery after nerve damage. Nature communications, 10(1), 1–14.

Sachdeva, R., Krassioukov, A. V., Bucksot, J. E., & Hays, S. A. (2020). Acute Cardiovascular Responses to Vagus Nerve Stimulation after Experimental Spinal Cord Injury. Journal of Neurotrauma, 37(9), 1149–1155.

Lewis, M. E., Al-Khalidi, A. H., Bonser, R. S., Clutton-Brock, T., Morton, D., Paterson, D., … & Coote, J. H. (2001). Vagus nerve stimulation decreases left ventricular contractility in vivo in the human and pig heart. The Journal of physiology, 534(Pt 2), 547.

Yamakawa, K., So, E. L., Rajendran, P. S., Hoang, J. D., Makkar, N., Mahajan, A., … & Vaseghi, M. (2014). Electrophysiological effects of right and left vagal nerve stimulation on the ventricular myocardium. American Journal of Physiology-Heart and Circulatory Physiology, 307(5), H722–H731.

Shinlapawittayatorn, K., Chinda, K., Palee, S., Surinkaew, S., Thunsiri, K., Weerateerangkul, P., … & Chattipakorn, N. (2013). Low-amplitude, left vagus nerve stimulation significantly attenuates ventricular dysfunction and infarct size through prevention of mitochondrial dysfunction during acute ischemia-reperfusion injury. Heart Rhythm, 10(11), 1700–1707.

Vimercati, C., Qanud, K., Ilsar, I., Mitacchione, G., Sarnari, R., Mania, D., … & Recchia, F. A. (2012). Acute vagal stimulation attenuates cardiac metabolic response to β-adrenergic stress. The Journal of Physiology, 590(23), 6065–6074.

Zhang, Y., & Mazgalev, T. N. (2009). Cardiac vagal stimulation eliminates detrimental tachycardia effects of dobutamine used for inotropic support. The Annals of thoracic surgery, 88(1), 117–122.

Berk, J. L., Hagen, J. F., Tong, R., & Maly, G. (1977). Pulmonary insufficiency produced by norepinephrine: a comparison with epinephrine. Circulatory Shock, 4(3), 247–251.

Ganzer, P. D., Colachis 4Th, S. C., Schwemmer, M. A., Friedenberg, D. A., Dunlap, C. F., Swiftney, C. E., … & Sharma, G. (2020). Restoring the Sense of Touch Using a Sensorimotor Demultiplexing Neural Interface. Cell. 181(4):763-773.e12. doi10.1016/j.cell.2020.03.054

Bouton, C. E., Shaikhouni, A., Annetta, N. V., Bockbrader, M. A., Friedenberg, D. A., Nielson, D. M., … & Morgan, A. G. (2016). Restoring cortical control of functional movement in a human with quadriplegia. Nature, 533(7602), 247–250.

